# Structure-based phylogenetic analysis reveals multiple events of convergent evolution of cysteine-rich antimicrobial peptides in legume-rhizobium symbiosis

**DOI:** 10.1101/2025.09.09.675119

**Authors:** Amira Boukherissa, Siva Sankari, Tatiana Timchenko, Mickaël Bourge, Peter Mergaert, George C. diCenzo, Jacqui A. Shykoff, Benoît Alunni, Ricardo C. Rodríguez de la Vega

## Abstract

Nitrogen is essential for plant growth, yet its availability often limits agricultural productivity. Some legumes have evolved a unique ability to form symbiotic relationships with nitrogen-fixing soil bacteria called rhizobia, enabling them to thrive in nitrogen-deficient soils. In five legume clades, an exploitive strategy has evolved in which rhizobia undergo Terminal Bacteroid Differentiation (TBD), where the bacteria become larger, polyploid, and have a permeabilized membrane. Terminally differentiated bacteria are associated with higher N_2_-fixation and, thus, a higher return on investment to the plant. In several members of the IRLC (Inverted Repeat-Lacking Clade) and the Dalbergioid clades of legumes, this differentiation process is triggered by a set of apparently unrelated plant antimicrobial peptides with membrane-damaging activity, known as Nodule-specific Cysteine-Rich (NCR) peptides. However, whether NCR peptides are also implicated in symbiotic TBD in other legume clades and whether they are evolutionarily related remains unknown. Here, to address the molecular identity of NCR peptides and their evolution in different legume clades, we performed inter- and intra-clade comparisons of NCR peptides in representative species of four TBD-inducing legume clades. First, we collected genomic and proteomic data of species for which NCR peptides are known (1523 NCR peptides). We then used sequence similarity-based clustering to regroup the NCR peptides, resulting in over 400 different NCR clusters, each clade-specific. We obtained Hidden Markov Models for each cluster and used them to predict NCR peptides in 21 legume genomes (6 clades), including newly generated deep-sequenced root and nodule RNA-seq data of *Indigofera argentea* (Indigoferoid clade) and newly assembled high-quality transcriptomes of *Lupinus luteus* and *Lupinus mariae-josephae* (Genistoid clade), using tailored gene prediction pipeline and transcriptome matching. This resulted in 3710 NCR peptides in species that induce TBD. To date, the rapid diversification of NCR peptides that reduces the sequence similarities has masked the origin of NCR peptide evolution. We obtained high-confidence structural models for one sequence of each cluster. We performed structure-based clustering and phylogenetics, which resulted in 23 superclusters (14 inter-clade and nine clade-specific) that we represent in a structural distance-based tree. Our study revealed that the evolution of NCR peptides is a mix of divergent and convergent processes within each clade. We further chose nine independently evolved NCR peptides to test *in vitro* whether they are functional analogs in symbiosis.

**Graphical abstract:** Overview of the experimental and computational workflow for NCR peptide detection, characterization, and structural analysis.
Nodule and root samples from *Indigofera argentea* (8 weeks post-inoculation) were collected and subjected to RNA extraction, library preparation, and Illumina PE150 sequencing. Raw RNA-seq reads from two *Lupinus* species were also included (*Lupinus luteus* and *Lupinus mariae-josephae)*. Bacteroid differentiation of *I. argentea* was assessed by flow cytometry and confocal microscopy. Transcriptomes were assembled de novo and analyzed for differential gene expression between root and nodule tissues. NCR peptides were identified from them and other legume genomes and transcriptomes using the SPADA pipeline and HMM profiles from NCR clusters of the known NCR peptides. The putative NCR peptides were filtered based on conserved cysteine motifs, length, and nodule expression to build an exhaustive NCR peptide database. 3D structural predictions of NCR clusters were performed using AlphaFold2 (pLDDT >70), followed by structural clustering (Foldseek) and phylogenetic analysis (Foldtree). Functional validation involved flow cytometry and antimicrobial assays (against *Eschericha coli*, *Sinorhizobium meliloti*, and *Bacillus subtilis*), enabling structural and evolutionary characterization of NCR peptides. The green box at the top represents the experimental analysis, the blue box represents the sequence-based computational pipeline, the red box represents the structure-based computational pipeline, and the grey box at the bottom left represents the functional validation and interpretation of the results.

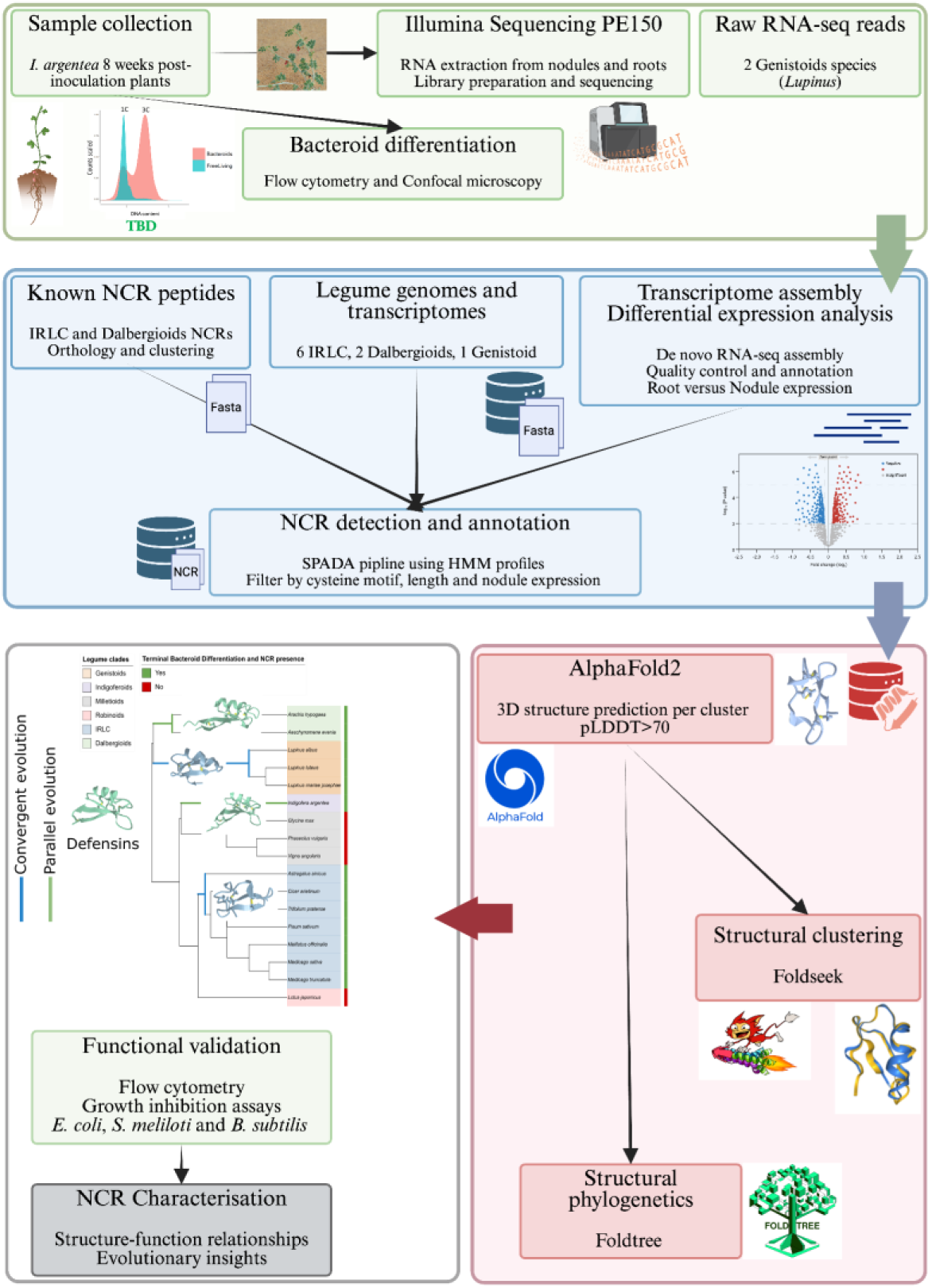

## INTRODUCTION

Legume plants (Fabaceae) have evolved the ability to house symbiotic nitrogen-fixing bacteria in their root nodules. When nitrogen is limited, legume plants can enter a symbiotic interaction with N_2_-fixing soil bacteria called rhizobia, an umbrella term that includes members of the classes *Alphaproteobacteria* and *Betaproteobacteria*. During this interaction, the legume plant forms root nodules where rhizobia are housed intracellularly as structures called bacteroids that fix atmospheric nitrogen and transfer ammonia to the plant. In return, legume plants provide these microsymbionts with carbon and other nutrients. This interaction initiates after mutual recognition between the host plant and a compatible bacterial partner, involving an exchange of signaling molecules between the two partners (Oldroyd, 2013). First, when grown in nitrogen-deprived soil, the legume plant releases flavonoids to the soil, thereby attracting rhizobia and triggering the bacteria’s production of nodulation (Nod) factors. On perceiving these Nod factors, the plant root hairs curl to trap the rhizobia and guide them, via infection threads, toward the incipient nodule. When they reach the cortical cells (Gage, 2004), the rhizobia are released by the infection threads and internalized by the nodule cells. Inside the nodule and, more precisely, inside the subcellular compartment called the symbiosome, rhizobial metabolism is rewired, and the rhizobia become nitrogen-fixing bacteroids.

Depending on the plant host, bacteroids may remain similar to free-living bacteria, with their shape and genome copy number unaltered during the symbiosis (Lamouche et al., 2019; Oono et al., 2010). However, in some legume plants like *Medicago truncatula* and relatives belonging to the Inverted Repeat Lacking Clade (IRLC), bacteroids undergo a process called Terminal Bacteroid Differentiation (TBD) (Haag & Mergaert, 2020; Lamouche et al., 2019; Oono & Denison, 2010). Terminally differentiated bacteroids are larger, elongated, or spherical cells with permeabilized membranes. They undergo endoreduplication and become polyploid while losing their ability to divide, but fix N_2_ more efficiently, providing a higher return on investment for the plant (Alunni & Gourion, 2016; Mergaert et al., 2006). In *Medicago* and its relatives, this differentiation process is induced by small plant antimicrobial peptides called NCR (Nodule-specific Cysteine-Rich) peptides, which are highly expressed in the nodules of some legume species that trigger TBD (Pan & Wang, 2017; Van de Velde et al., 2010). These peptides are composed of a signal peptide, which drives their secretion, and a 20 to 50-amino acid-long mature peptide, including 4, 6, or 8 cysteines that form two, three, or four disulfide bridges (Haag et al., 2012; Montiel et al., 2017; Ribeiro et al., 2017; Shabab et al., 2016). The mature peptides are highly variable at the sequence level except for the cysteine pattern (Mergaert et al., 2003). The number of NCR peptides among legume species varies from 7 (*Glycyrrhiza uralensis*) to 700 (*Medicago truncatula*) (Montiel et al., 2017; Young et al., 2011). According to their isoelectric point, NCR peptides can be classified as cationic, neutral, or anionic. In the few cases studied, cationic NCR peptides display antimicrobial activity by permeabilizing bacterial membranes *in vitro*, while no activity has been shown for anionic NCR peptides (Maróti et al., 2011, 2015) Furthermore, NCR peptides have been suggested to provoke a cell cycle switch in symbiosis, where it has been demonstrated that NCR247 can inhibit bacterial cell division by interacting with the FtsZ protein (Farkas et al., 2014). Moreover, it has been shown that NCR247 interacts with other bacterial proteins, such as ribosomal proteins and the chaperonin GroEL (Farkas et al., 2014).

Nodule Cysteine-Rich peptides are required for terminal bacteroid differentiation and establishing an effective symbiosis between IRLC legumes and rhizobia (Van de Velde et al., 2010), where they control the bacterial life cycle and other cellular pathways (Roy et al., 2020). These peptides are expressed in waves during different stages of nodule formation and bacteroid differentiation (Guefrachi et al., 2014). Recently, it has been shown that at least nine individual NCR peptides are essential for the symbiosis in *Medicago truncatula* (Horváth et al., 2015, 2023; Kim et al., 2015). Furthermore, it has been demonstrated recently that NCR247 is involved in iron homeostasis, interacting with heme moieties and thereby facilitating iron uptake by rhizobia (Sankari et al., 2022).

Rhizobia can tolerate the stress provoked by NCR peptides and prevent membrane damage with the help of specific ABC transporters called BacA or BclA (Glazebrook et al., 1993), which are essential for effective symbiosis involving legumes that trigger TBD (Guefrachi et al., 2015; Haag et al., 2011). Mutants with deletion of the *bacA* gene cannot transport NCR peptides and die in the presence of cationic NCR peptides (Barrière et al., 2017).

TBD has been observed in five different legume clades (Genistoids, Mirbelioids, IRLC, Millettioids (= Indigoferoid/Millettioid), and Dalbergioids) (Oono et al., 2010). The role of plant-secreted NCR peptides in this process is known only in two of them, IRLC and Dalbergioids (Czernic et al., 2015; Montiel et al., 2017). NCR peptides from the IRLC and Dalbergioid clades have different sequences and cysteine motifs, but both induce TBD in the symbiont (**Figure 1**) (Czernic et al., 2015). Indeed, NCR peptides may have evolved independently in IRLC and Dalbergioid clades, supporting the idea of convergent evolution driving symbiont terminal differentiation (**Figure 1**) (Downie & Kondorosi, 2021). Nevertheless, a recent phylogenetic study between plant defensins and NCR peptides demonstrated that they may share the same origin (Salgado et al., 2022). However, all past studies about NCR peptides were sequence-based and limited to two clades and a small subset of NCR peptides. Therefore, the presence of NCR peptides in other clades, their molecular identity, and their evolution remain unknown.

**Figure 1.**
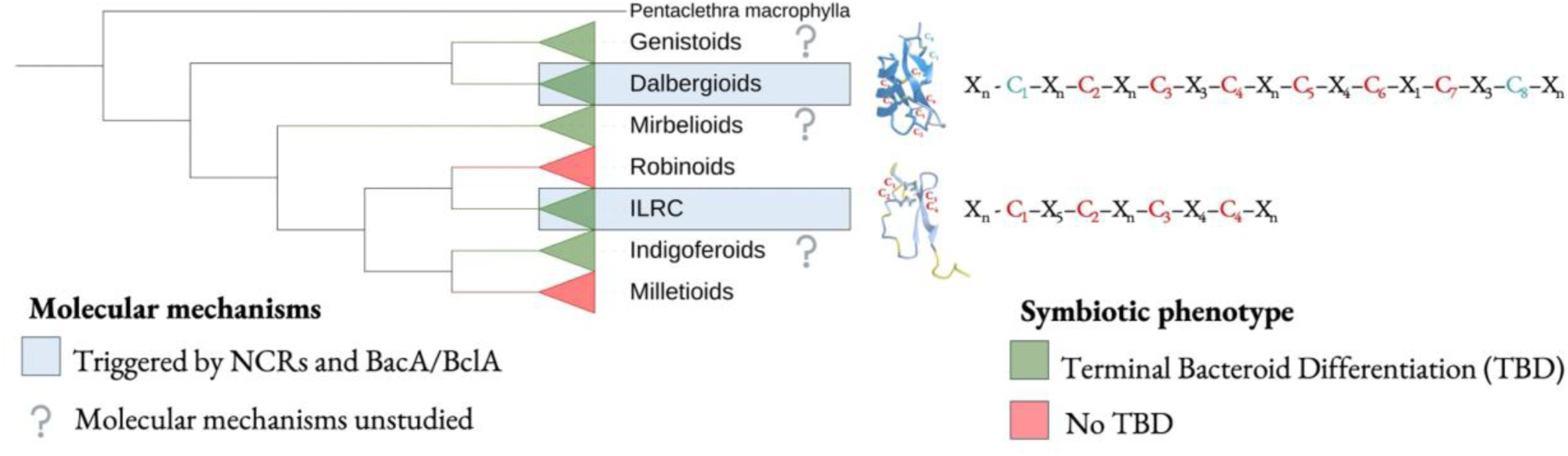
The evolution of terminal bacteroid differentiation (TBD) and NCR peptide mechanisms across legumes. Simplified legume phylogeny showing clades that undergo TBD (green) or do not (red). The blue shading highlights lineages where NCR peptides and BacA/BclA have been identified as triggers of TBD, and question marks represent species that undergo TBD with unstudied molecular mechanisms. Schematic NCR cysteine motifs and representative 3D structures illustrate the cysteine-rich β-sheet fold of NCR peptides, with variable sequences and loop regions. This figure introduces the central question of this study: **Do all cases of TBD involve NCR peptides, and how have these peptides evolved across legumes?**

The evolutionary analysis of protein families was traditionally focused on sequence-based approaches, using tools like BLAST (Altschul et al., 1990) for similarity searches, Hidden Markov Models (HMM) (Eddy, 1998) or profile-based detection, and multiple sequence alignments for phylogenetic reconstruction. While these methods have proven valuable for studying conserved protein families, they face significant limitations when applied to small and rapidly evolving peptides like NCR peptides. The high divergence observed in NCR peptides, with sequence identity often below 20%, even within the same clade (Alunni & Gourion, 2016; Montiel et al., 2017), hides potential evolutionary relationships and can lead to incomplete conclusions about their origins. This sequence-based limitation needs other approaches to uncover the true evolutionary history of these functionally essential peptides.

Structural biology offers a powerful complementary approach, as protein tertiary structure is generally more conserved than primary sequence during evolution (Illergård et al., 2009). Nevertheless, experimental structure determination through X-ray crystallography, nuclear magnetic resonance spectroscopy, or cryo-electron microscopy remains costly and time-consuming, particularly for large-scale comparative studies involving thousands of peptides. The recent revolution in computational structural biology, particularly through AlphaFold2’s deep learning-based structure prediction (Jumper et al., 2021), has resulted in the unprecedented accessibility of high-quality protein structures. This groundbreaking advancement enabled the emergence of structure-based clustering and phylogenetic approaches at an unparalleled scale, allowing researchers to integrate tools like Foldseek (van Kempen et al., 2024) for rapid structural clustering and Foldtree (Moi et al., 2023) for structure-based phylogenetic reconstruction. In the context of NCR peptides, where sequence similarity searches fail to reveal inter-clade relationships, structural analysis becomes valuable and essential for understanding their evolutionary origins and functional convergence across different legume lineages.

Here, we examine how NCR peptides evolved in legumes and how they are associated with TBD (**Figure 1**). We report an exhaustive list of NCR peptides of four legume clades that undergo TBD, grouped by sequence-based clusters and structure-based superclusters. Although the sequence analysis demonstrates that NCR peptides are legume clade-specific and highly diverse, the structural analysis grouped NCR peptides from different legume clades together and detected a hundred clusters in the same structural supercluster. This suggests that the NCR peptides of different legume clades may have evolved from multiple genes of the same gene family. Moreover, the presence of NCR-like genes in legume clades that do not induce TBD further suggests that NCR genes evolved from the same gene family but gained the function of inducing TBD independently in some legume clades. This paper highlights the importance of structure-based analysis in studying the evolution of protein families with highly diverse sequences.

## RESULTS

### NCR peptides are clade-specific at the sequence level

To decipher the evolutionary history of NCR peptides, we first studied the known NCR peptides from IRLC and Dalbergioid clades at intra-clade and inter-clade scales. We performed homology, orthology, and Markov clustering analysis of all proteins (including NCR peptides) of all legume species where NCR peptides are known and the inferred proteomes are available, i.e., *Medicago truncatula*, *Medicago sativa, Cicer arietinum,* and *Pisum sativum* from IRLC, and *Arachis hypogaea* and *Aeschynomene evenia* from Dalbergioids (see Materials and Methods). This approach led us to identify 63,490 orthologous clusters from 483,710 proteins. Among those clusters, 18,856 were inter-clade (containing proteins from both IRLC and Dalbergioid clades), and 44,634 were clade-specific (36,792 IRLC clusters and 7,842 Dalbergioid clusters). In this initial analysis, all NCR peptide clusters are legume clade-specific **(Figure S1)**. In the IRLC, 1492 of the 1523 NCR peptides were clustered (assigned to an orthologous group) into 651 clusters. Of them, 203 clusters exclusively contain NCR peptides (568 NCR peptides), and the remaining 448 clusters (924 NCR peptides) were NCR-mixed orthologous groups with at least one NCR peptide and one non-NCR protein. Among the 448 NCR-mixed clusters, 238 clusters were NCR-monotypic clusters with only one NCR peptide in the cluster. In the Dalbergioid clade, 117 of the 155 NCR peptides were clustered into 20 clusters. Of them, 7 clusters exclusively contain NCR peptides (40 NCRs), and the remaining 13 clusters (77 NCRs) were NCR-mixed orthologous groups, seven of which were NCR-monotypic. One big mixed cluster with 53 sequences represented almost all the *Aeschynomene evenia* NCR peptides. In contrast, the other clusters were small clusters of *Arachis hypogaea* NCR peptides, highlighting the between-species sequence divergence of NCR peptides even within a clade. Furthermore, all the IRLC NCR peptide clusters contain only NCR peptides with four or six cysteine motifs (hereafter called type-1) in the mature peptides **(Figure 2d)**, while 95% of the NCR peptides in the Dalbergioids NCR clusters had a defensin-motif with eight cysteines (type-2) **(Figure 2d)**. For further analysis, we considered only the NCR peptides of clusters (including NCR-mixed clusters) containing at least two NCR peptides after filtering out the NCR peptide sequences without a signal peptide. Consequently, we end up with 385 IRLC NCR clusters (1191 NCRs) and 11 Dalbergioids NCR clusters (102 NCRs). Together, these results revealed a clear clade-specific organization and diversification of NCR peptides, providing a robust foundation for understanding their evolutionary mechanisms across legume species.

**Figure 2.**
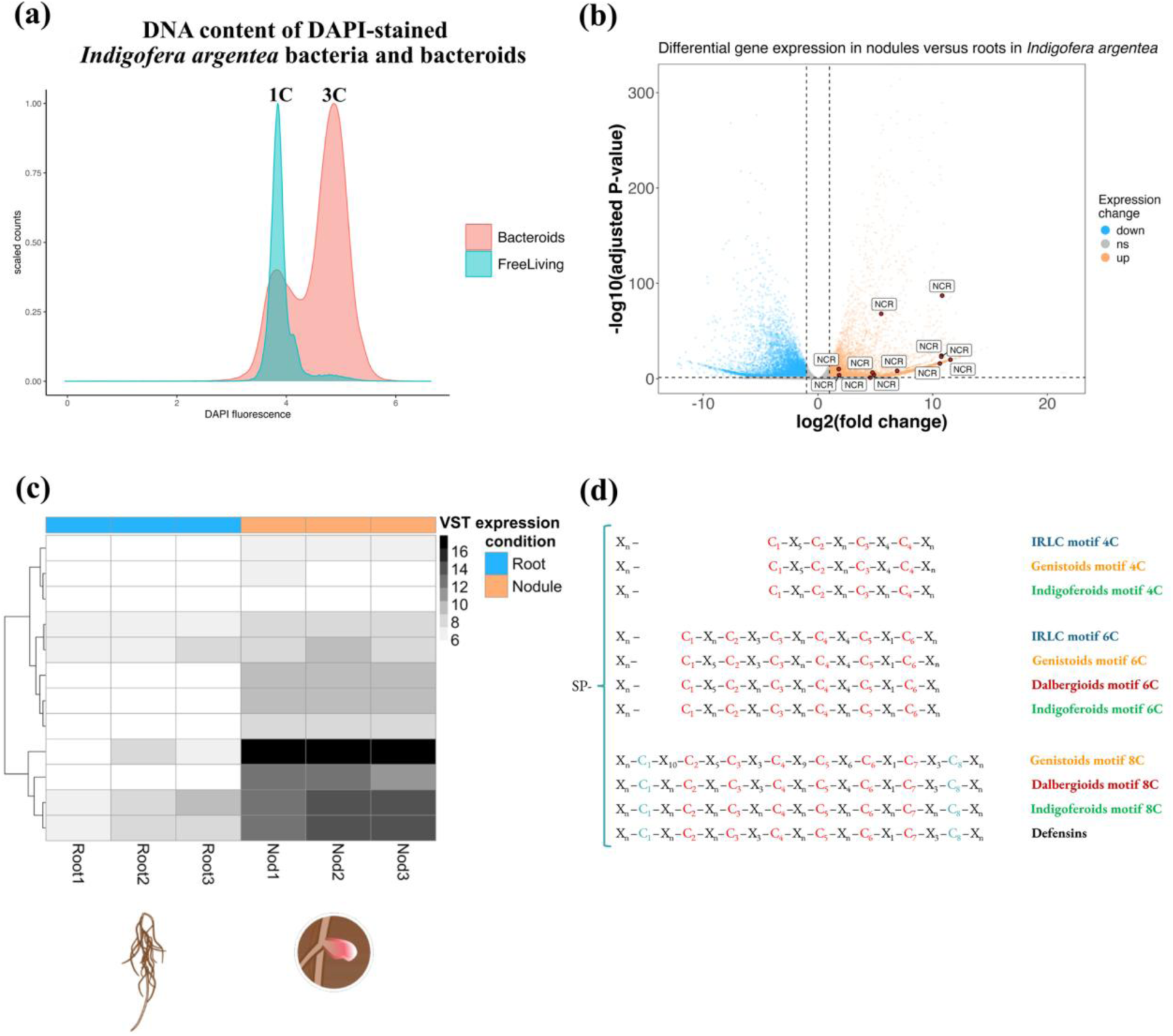
*Indigofera argentea* induces TBD and expresses NCR peptides. The few differentially expressed NCR peptides in nodules of *I. argentea* may induce moderate TBD with only 3C. (**a**) The DNA content of DAPI-stained *B. elkanii* strain SA281 bacteria and bacteroids isolated from *I. argentea* nodules was measured by flow cytometry, and (**b**) Volcano plot of all *I. argentea* transcripts where the down-regulated transcripts (Log2FoldChange < -1) in nodules are in blue (left), the up-regulated transcripts (Log2FoldChange > 1) are in orange (right), the transcripts that had non-significant expression difference between roots and nodules are in gray (middle and bottom) and the *NCR* genes among the up-regulated transcripts are annotated and colored in red. (**c**) Heatmap of the expression of 12 differentially expressed NCR genes in *I. argentea* nodules (blue) and roots (orange), each with three biological replicates. Gene expression values were transformed using DESeq2’s variance stabilizing transformation (VST) to reduce mean-dependent variance and accurately compare samples. Expression values were centered and scaled per gene. The color scale represents VST-transformed (Variance Stabilizing Transformation) expression values, ranging from 0 (white) to 16 (black), with higher values indicating higher expression. (**d**) Conserved cysteine motifs of NCR-like peptides across legume clades. Motifs are shown as “Cn” for conserved cysteine residues and “Xn” for stretches of n amino acids, where n corresponds to the most frequent spacing observed in the sequences. The red cysteines represent NCR-motif (type-1 in this paper), and blue cysteines represent defensin-motif (type-2).

### Sequence-based clusters help identify new NCR peptides in understudied legume clades

In addition to IRLC and Dalbergioids, some Indigoferoids and Genistoids induce TBD (Oono & Denison, 2010), but whether NCR peptides are also involved remains unknown. Therefore, to expand our NCR peptide dataset to other legume clades, we generated deep-sequenced root and nodule RNA-seq data for *Indigofera argentea*, a legume from the Indigoferoid clade. We first confirmed that *I. argentea* induces TBD, for which we quantified the DNA content and size of the bacteroids in the nodules of *I. argentea* using flow cytometry. This analysis showed an increase in the DNA content with peaks at 3C, albeit with only slightly enlarged bacteroids (**Figure 2a; Figure S2**). We obtained a *de novo* reference transcriptome assembly and annotation using the generated *I. argentea* nodule and root RNA-seq data, and we also assembled and annotated publicly available raw RNA-seq datasets sequenced from the nodules and roots of two *Lupinus* species from the Genistoid clade (*L. luteus* and *L. mariae-josephae*) (Keller et al., 2018). We performed differential expression analysis of nodules and roots.

The *de novo* assembly of the *I. argentea, L. luteus,* and *L. mariae-josephae* transcriptomes generated 277,022, 156,834, and 152,943 contigs, respectively. Assembled contigs include different splice variants of one gene, which we subsequently merged into one contig called a “supertranscript”. The resulting assemblies contain 72,846, 57,642, and 55,700 supertranscripts for *I. argentea, L. luteus,* and *L. mariae-josephae,* respectively. Based on BUSCO genome completeness and the mapping of reads against the de novo assemblies’ metrics, the assemblies are of high quality, with more than 99% of the cleaned reads mapped to their corresponding contigs, and >85% could be mapped to their corresponding “supertranscript” (**Dataset S2**). Moreover, 95% of the Viridiplantae and 86% of the Fabales BUSCO genes were identified as complete and single-copy for *I. argentea* (**Figure S3**).

We next used a hidden Markov model-based gene caller, SPADA, to search for NCR peptides in the de novo transcriptome assemblies of these three species (*I. argentea, L. luteus,* and *L. mariae-josephae*) and the published genome of the related organism *Lupinus albus*. SPADA was run once for the three Genistoids species and run twice for *I. argentea* (see below). A total of 129, 238, 259, and 398 putative NCR peptides were identified in *I. argentea, L. luteus, L. mariae-josephae,* and *L. albus*, respectively. Following this analysis, 12, 87, 69, and 36, respectively, were confirmed as “NCR peptides” due to their differential expression in nodules and the presence of at least four cysteines in the predicted mature peptide. A maximum of 100 amino acids was allowed in the initial SPADA run. However, some predicted NCR peptides were longer than those reported in other species under this filter. Consequently, the length criterion was relaxed in the subsequent run to identify additional candidates. The mature NCR peptides’ length ranged from 23 to 85 in *L. luteus*, 23 to 95 in *L. mariae-josephae*, and 23 to 65 in *L. albus* (**Figure 3a**). However, in *I. argentea,* the length range of the mature NCR peptide was from 46 to 152, which is higher than expected (**Figure 3a**). Half of the *I. argentea* NCR peptides had an NCR motif with four or six cysteines, and the other half had a defensin motif with eight cysteines. For the *Lupinus* spp., only around 14% of the annotated NCR peptides had a defensin motif.

**Figure 3.**
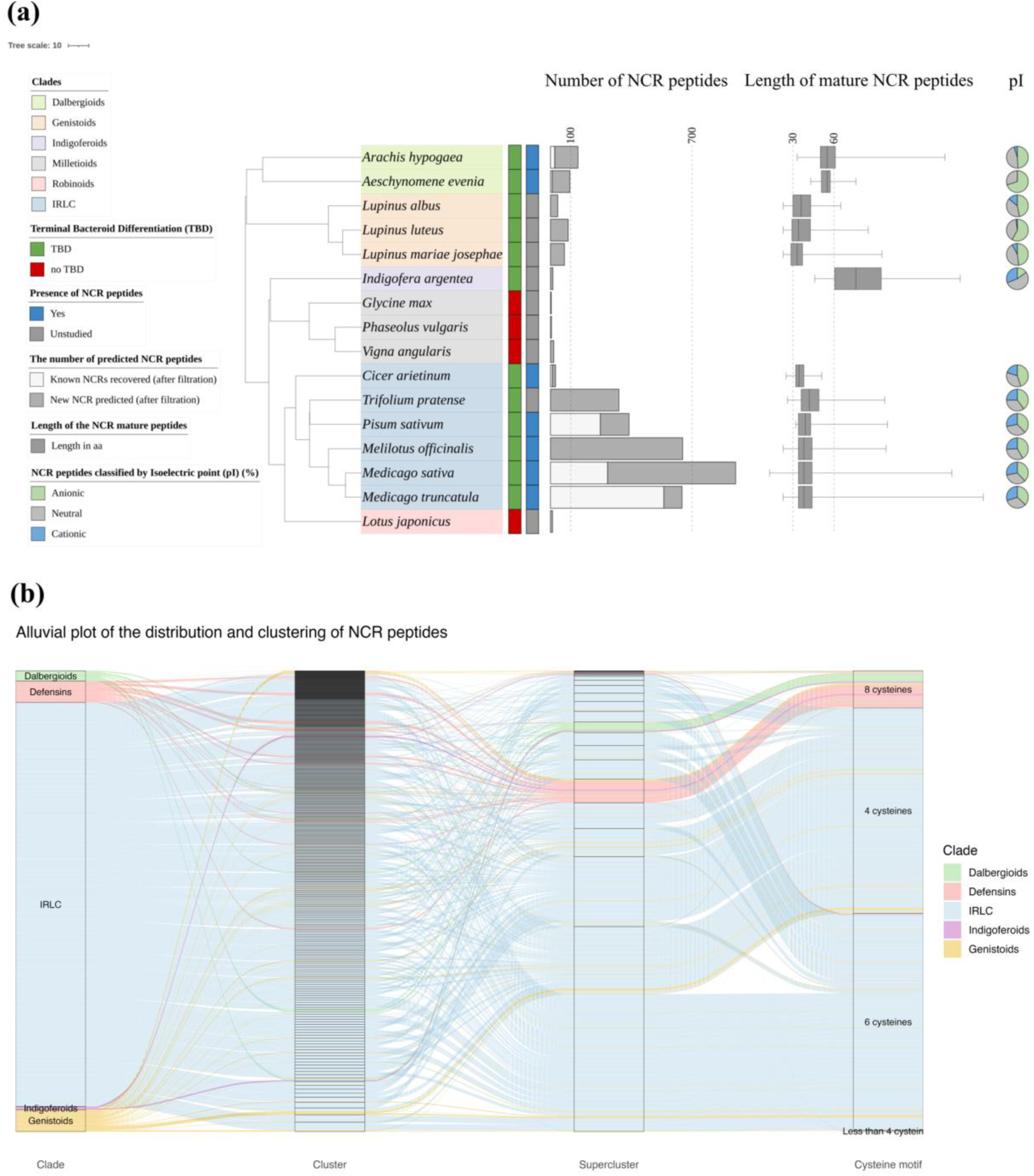
The distribution and characteristics of NCR peptides across the legume phylogeny, NCR clusters, and superclusters. (**a**) Legume species tree generated by TimeTree (Kumar et al., 2017), where the labels are the species colored by clade (top left legend). The first strip represents the occurrence of TBD (green) or not (red) (second legend from the top). The second strip represents the presence (blue) or absence (grey) of NCR peptides identified according to previous studies (third legend from the top). The bar plots represent the number of recovered known NCR peptides (white) and the new NCR peptides from the predicted NCR peptides after filtration (grey) (the fourth legend from the top). The box plots represent the length of the mature peptides (the fifth legend from the top), and the pie chart plots represent the percentage of anionic (pI <= 6) (green), cationic (pI >= 8) (blue), and neutral (6 < pI < 8) (grey) peptides (the last legend from the top). (**b**) Alluvial (RiverPlot) of all NCR peptides regrouped from the left by clades, sequence-based clusters, structure-based superclusters, and cysteine motifs, where we can trace the fate of each NCR peptide or cluster. For instance, IRLC NCR peptides regroup with Genistoids in the same clusters and superclusters, while Dalbergioids and Indigoferoids regroup with defensins in the same superclusters.

The differential expression analysis showed 13,540 up-regulated and 12,356 down-regulated genes in the nodules of *I. argentea* compared to the roots (**Figure S4**). The 12 NCR peptides in *I. argentea* are highly and differentially expressed in nodules (**Figure 2b,c**), one of which is among the 10 most abundant transcripts. Additionally, three of the four NCR peptides most differentially expressed in nodules (clustering together in the heatmap) (**Figure 2c**) are cationic. Furthermore, the transcript abundances calculated using TPM (Transcripts Per Million) in the three *Lupinus* species show a significantly lower expression of NCR peptides in *Lupinus albus* than in the two other *Lupinus* species (**Figure S5**). In addition, *L. albus* has fewer NCR peptides than the two other *Lupinus* spp. (**Figure 3a**).

We classified the newly identified NCR peptides based on the Hidden Markov Models (HMM) constructed from multiple sequence alignments of the 396 sequence-based IRLC and Dalbergioid clusters (385 and 11 clusters, respectively). We found no matching profile for 11 of the 12 *I. argentea* NCR peptides; thus, at the sequence level, *I. argentea* NCRs are also largely clade-specific. In contrast, 75 out of 87 *L. luteus*, 61 out of 69 *L. mariae-josephae*, and 24 out of 36 *L. albus* NCR peptides matched HMM profiles of the IRLC clusters, with one additional *L. luteus* sequence matched to a HMM profile of a Dalbergioid NCR cluster. This leaves 11, 8, and 12 of the *L. luteus*, *L. mariae-josephae*, and *L. albus* NCR peptides as clade-specific at the sequence level, respectively. According to this sequence-based approach, none of the NCR peptides previously identified as necessary in the *Medicago truncatula* - *Sinorhizobium meliloti* symbiosis (NCR247, NCR211, NCR169, NCR-new35, NCR343, NCR086, NCR314, NFS1, and NFS2) (Horváth et al., 2015, 2023; Saifi et al., 2024; Wang et al., 2017; Yang et al., 2017) would have homologs in *I. argentea, L. luteus*, *L. mariae-josephae*, or *L. albus.* In summary, these results reveal novel NCR peptides in Indigoferoids and Genistoids legumes, with *I. argentea* encoding largely clade-specific variants, while most *Lupinus* NCR peptides share sequence similarity with previously characterized NCR peptides from IRLC, leading to the first described inter-clade clusters (**Figure S1**).

### Legume-wide distribution of NCR peptides across legume species

Once our dataset was expanded to other legume clades, we also expanded our NCR peptide dataset inside each clade and each species. Specifically, we searched for new NCR peptides in all legume species where genomic and nodule transcriptomic data were available (14 in total with *Lupinus albus*, **Dataset S1**) using SPADA that took as input the HMM profiles built from our IRLC and Dalbergioids NCR clusters and the CRP (Cysteine-Rich Peptide) clusters (a broader family of small, cysteine-rich peptides that includes defensins and other antimicrobial peptides) (Nallu et al., 2014) (see Materials and Methods). This analysis allowed us to recover NCR peptides in 14 legume species, including four species for which NCR peptide repertoires have never been described (**Dataset S3**) and novel NCR peptides in well-studied clades (e.g., 13% to 70% of the NCR peptides in six IRLC species were newly identified here) (**Figure 3a**). While almost all newly identified NCR peptides in the IRLC and Genistoids were classified into our existing NCR clusters, in the Dalbergioids, only 24 to 32% of the NCR peptides were classified, and only one of the recovered Indigoferoid NCR peptides (by the second SPADA run) was classified into one Dalbergioid cluster (**Dataset S3**). Additionally, the six *I. argentea* NCR peptides recovered in the first SPADA run match a profile constructed from CRPs. A second round of SPADA, using HMM profiles built from our expanded NCR peptide clusters (including the newly identified NCR peptides from IRLC, Dalbergioids, and Genistoids) and supplemented with one *I. argentea* NCR peptide HMM from (Ren, 2018), recovered six new *I. argentea* NCR peptides, five matching the *I. argentea* profile and one matching a profile from a Dalbergioid cluster.

The average length of the mature NCR peptides was 34 for IRLC and 37 for Genistoids. The mean lengths of the Dalbergioids and Indigoferoids mature NCRs were 55 and 81 aa, respectively (**Figure 3a**). As expected (Zorin et al., 2022), the anionic NCR peptides are the most abundant in all the studied legume species, except for *I. argentea*, where the neutral NCRs are the most abundant (**Figure 3a**). *Medicago* spp. from the IRLC clade had the highest percentage of cationic NCR peptides (36%), while *Aeschynomene evenia* from the Dalbergioids clade had the lowest rate with no cationic NCR peptides (**Figure 3a**). More generally, the Genistoids and the Dalbergioids clades had few to no cationic NCR peptides. As TBD has been observed in both clades, this suggests that cationic NCR are not strictly necessary to induce bacteroid formation.

Furthermore, to reveal the evolutionary connections between NCR clusters, we performed a sequence similarity network analysis using CLANS (CLuster ANalysis of Sequences), a tool that visualizes sequence relationships in 3D space based on pairwise BLAST similarities. In CLANS, sequences with significant similarity are connected by lines and cluster together through force-directed graph layout, creating a visual map of evolutionary relationships. Our analysis of representative sequences from each NCR peptide cluster revealed three distinct groups in the network (**Figure 4a**). The IRLC and Genistoids NCR peptides formed an extensive, interconnected network at the center, indicating their close evolutionary relationship despite originating from different clades. In contrast, the Dalbergioids NCR peptides formed a separate, more dispersed cluster, suggesting greater sequence divergence within this clade. The defensins formed a third, tightly clustered group (**Figure 4a**). The Indigoferoid and a few *Lupinus* NCR peptides appeared as small, isolated clusters at the network’s periphery (**Figure 4a**). Interestingly, one Dalbergioid cluster showed unexpected proximity to the defensin group (**Figure 4a**), suggesting a potential evolutionary relationship. However, the overall divergence of amino acid sequences between NCR peptides from different clades indicates that rapid sequence evolution may mask some evolutionary connections.

**Figure 4.**
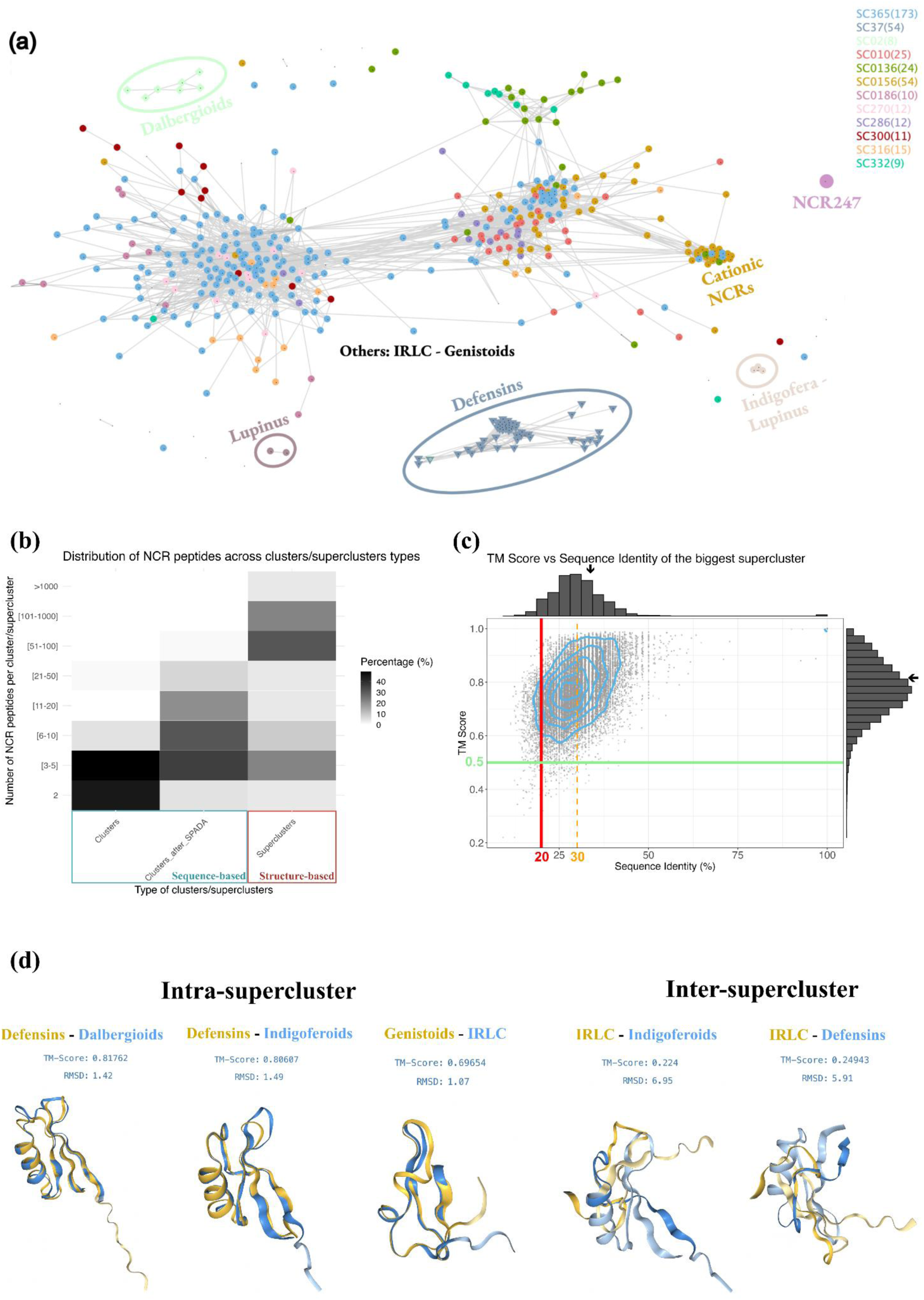
Structural conservation of NCR peptides displaying a high level of sequence divergence. (**a**) CLANS (Frickey & Lupas, 2004) sequence similarity network of the identical sequences used to perform structural analysis (one per cluster + monotypic). The colors are the same as used to represent the superclusters in the structure-based phylogeny of Figure 4. (**b**) Heatmap of the percentage of NCR peptides clustering according to the number of NCR peptides in the cluster or supercluster. Small clusters are abundant when we consider only the IRLC and Dalbergioids NCR peptides identified in the initial analysis (clade-specific), whereas cluster size is moderate when we consider the additional NCR peptides identified in the expanded analysis that also includes the Indigoferoids and Genistoids NCR peptides, and large for superclusters produced by classifying the NCR peptides by their structures. (**c**) Dot plots and histograms that represent the sequence identity between sequences and the TM scores between structures of NCR peptides of the biggest supercluster, where we demonstrated that inside one supercluster, the sequences are divergent (sequence identity around 27%), but their structures are conserved (TM score around 0.75). The two arrows in the sequence identity and TM score distributions represent the median. The horizontal green line represents the threshold of the TM score, where structures with TM scores above 0.5 are considered similar. The vertical red line represents the expected sequence identity of two random proteins (<20%), and the vertical orange line represents the threshold below which homology is uncertain (30%, so-called the twilight zone). (**d**) Structural alignments of intra-supercluster structures with high TM scores versus inter-superclusters with low TM scores.

In summary, this expanded analysis increased the number of NCR peptide clusters from approximately 1,500 to more than 3,500 (**Figure 3b**), revealed NCR peptide clusters with a greater number of sequences per cluster (**Figure 4b**), and demonstrated that NCR peptide clusters are no longer clade-specific (**Figure S1**), reflecting a substantial diversification and inter-clade convergence of NCR peptides across the legume phylogeny.

### Structure-based analysis reveals the hidden shared evolutionary history of NCR peptides

NCR peptides are small (30-50 aa in the mature peptide, with few exceptions) and are divergent at the sequence level. This rapid diversification of NCR peptides that reduce the sequence similarities has likely hidden the evolutionary origin of these peptides and made the inference of their evolutionary history using traditional phylogenetic analysis very difficult. Therefore, to gain more insights into the evolution of NCR peptides, we used structural clustering and phylogenetics to study the NCR peptides at the tertiary structure level, to see whether they are also divergent at the structural level and to reduce the number of clusters, regrouping them into structural superclusters.

To perform structural analysis, we predicted the tertiary structures of one representative NCR peptide from each of 396 clusters (see Materials and Methods) and 48 defensins (outgroup) from four legume clades that induce TBD using AlphaFold2 (Jumper et al., 2021). After filtering out the structures with a pLDDT (predicted Local Distance Difference Test) score < 70, 390 NCRs and 48 defensins were kept. The tertiary structures of the unclassified clade-specific NCR peptides from *I. argentea* and the *Lupinus* species (Genistoids) were also predicted using AlphaFold2. We excluded five *I. argentea* NCR peptides as we could not predict their tertiary structure with a confident pLDDT score (pLDDT<55). We used Foldseek (van Kempen et al., 2024) to regroup all structures into 23 superclusters, nine of which were small legume clade-specific clusters and 12 of which were big inter-clade clusters (**Figure 4b**). For instance, this analysis allowed us to regroup the cationic clusters of IRLC and Genistoids in the same supercluster (SC156), including the NCR peptide NCR343 that was identified as essential for an effective symbiosis in *M. truncatula* (**Figure 5**) (Gao et al., 2023). Defensins group together in one supercluster with a few Genistoid NCR peptides, Dalbergioid NCR peptides, and one of the most abundant Indigoferoid NCR peptides (**Figure 5**). The well-studied peptide NCR247 from *M. truncatula* belongs to a depauperate supercluster composed of just five sequences, exclusively found in *Medicago* species.

**Figure 5.**
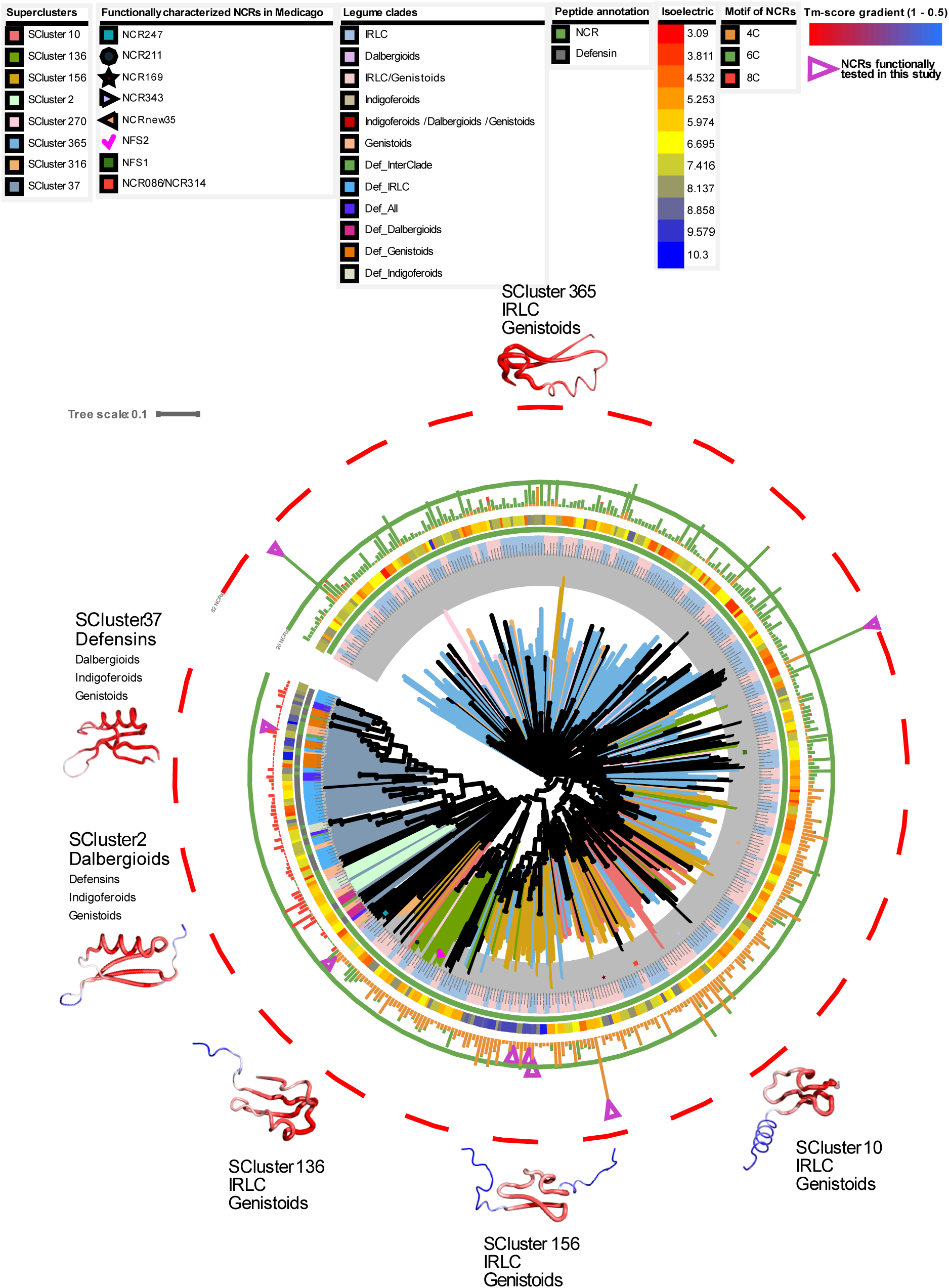
Structural phylogenetic analysis of NCR peptides across legumes. Phylogeny of 444 tertiary structures of NCR peptides (one per cluster) and defensins produced with Foldtree, where the branches are colored according to the superclusters defined with Foldseek (van Kempen et al., 2024). The identified NCR peptides important for effective symbiosis in *M. truncatula* in previous studies are annotated in the branches with different motifs. The labels represent the NCR clusters. The labels are colored according to the legume clades. In the first strip, starting from the inner strip, the red represents NCR peptides and the green defensins. The second strip represents the mean isoelectric point of the clusters, calculated from the mean isoelectric points of all NCR peptides of each cluster. The multibar plots represent the size of clusters (number of NCR peptides), where the red represents the number of NCR peptides with an 8-cysteine motif, green represents the number of NCR peptides with a 6-cysteine motif, and orange represents the number of NCR peptides with a 4-cysteine motif. The structural alignments represented by a sausage representation are the alignments of all the structures of the supercluster with a color gradient from red (TM score 1) to blue (TM score 0.5). The size of the names of the legume clades or defensins inside the superclusters is proportional to the number of NCR peptides from this clade in the supercluster.

The comparison between the TM (Template Modeling) scores (Y. Zhang & Skolnick, 2005) and the sequence identities inside the biggest supercluster (**Figure 4c**) of IRLC and Genistoids showed that the tertiary structures of NCR peptides are relatively conserved despite the divergence of their amino acid sequences, which can explain why the evolutionary connections were hidden using sequence-based approaches. Moreover, the TM scores within the Dalbergioids supercluster were generally higher than the TM scores between members of this supercluster and all the other superclusters (**Figure S7**). A different pattern was seen for a focal IRLC-Genistoids supercluster. In this case, there is one distribution representing comparisons between the focal IRLC-Genistoids supercluster and the Dalbergioids-Defensins superclusters and other IRLC-Genistoids superclusters (**Figure S7** in yellow), which overlapped with the other blue distribution that represents the distribution of TM scores for intra-supercluster comparisons within the focal IRLC-Genistoids supercluster.

### Structure-based phylogeny uncovers supercluster relationships and evolutionary history of NCR peptides

In order to decipher the evolution of NCR peptides and defensins and have a better overview of our superclusters, we used Foldtree (Moi et al., 2023), a structural phylogenetic approach that uses structural distances to build a tree (see Materials and Methods). This analysis allowed us to build structure-based trees for all our structures and each supercluster (**Figure 5**). These structure-based trees were supported by structural alignments of each supercluster (see Materials and Methods). The structural phylogenetic tree regrouped together the superclusters of the defensins and Dalbergioid NCR peptides, separately from the IRLC-Genistoid NCR peptide superclusters (**Figure 5**). Additionally, the structural alignment of the defensin supercluster is similar to the Dalbergioid NCR peptide supercluster, while both are different from the structural alignments of the IRLC-Genistoid NCR peptide superclusters. Albeit not fully resolved, two scenarios emerged from these results, revealing the hidden evolutionary history of NCR peptides. On one hand, NCR peptides from Dalbergioids and Indigoferoids appear to have evolved from defensins (**Figure 4d**, **5, S6**). On the other hand, taking into consideration that IRLC and Genistoids are relatively distant clades, the NCR peptides in these two clades appear to have been recruited independently by convergent evolution and were then expanded and rapidly diversified in the IRLC clade (**Figure 4d**, **5, S6**). Indeed, the NCR peptides in these two clades regroup in the same clusters and superclusters with some species-specific NCR peptides.

In summary, our integrative structural phylogenetics pipeline revealed that despite extreme sequence divergence, NCR peptides retain conserved tertiary structures within and across legume clades, enabling the recovery of evolutionary relationships previously hidden by rapid sequence evolution. Notably, the structural phylogeny (**Figure 5**), the structural alignments (**Figure 4d**, **5**), and supercluster organization support two distinct evolutionary histories, one where Dalbergioid and Indigoferoid NCR peptides appear derived from defensin-like ancestors, and another where IRLC and Genistoid NCR peptides represent a case of convergent evolution, independently recruited and diversified for symbiotic function.

### Convergent evolution of NCR peptides function across distinct structural superclusters

NCR247 is a well-characterized cationic NCR peptide from *M. truncatula* that exhibits potent antimicrobial activity against a range of gram-negative and gram-positive bacteria (Farkas et al., 2014; Guerra-Garcia & Sankari, 2025; Jenei et al., 2020; Montiel et al., 2017; Penterman et al., 2014; Sankari et al., 2022). It also induces symbiotically essential processes, including driving of polyploidy and heme binding. In contrast, the anionic and neutral NCR peptides tested from *M. truncatula* do not exhibit potent antibacterial activity, and their functions remain largely unexplored. To validate the approach used to predict the NCR peptides, most importantly, to validate that the NCR peptides that regroup with defensins are truly NCR peptides, we synthesized nine evolutionarily distant NCR peptides from different clades and different superclusters (**Figure 5**; **Table 1**) and compared their activity directly with that of NCR247. We selected two cationic *C. arietinum* (IRLC) NCR peptides from the biggest NCR peptide cluster belonging to the biggest supercluster, one predicted in this study, and one that was previously known. Additionally, we picked two cationic NCR peptides from the third biggest cluster, one from *Astragalus sinicus* and one from *T. pratense* (IRLC), three NCR peptides from a cationic supercluster: one *M. sativa*, one *M. truncatula*, and one *T. pratense* (IRLC). Lastly, we selected one highly expressed and cationic *I. argentea* (Indigoferoid) NCR peptide belonging to the defensins supercluster. In addition, one highly expressed neutral *L. luteus* (Genistoid) NCR peptide from the second-biggest cluster was also chosen to see if its activity matches the tested neutral peptides from previous studies. The mass of these peptides ranged from 3.23 to 5.75 kDa, and their pI ranged from 6.9 to 10.31 (**Table 1**).

**Table 1.**
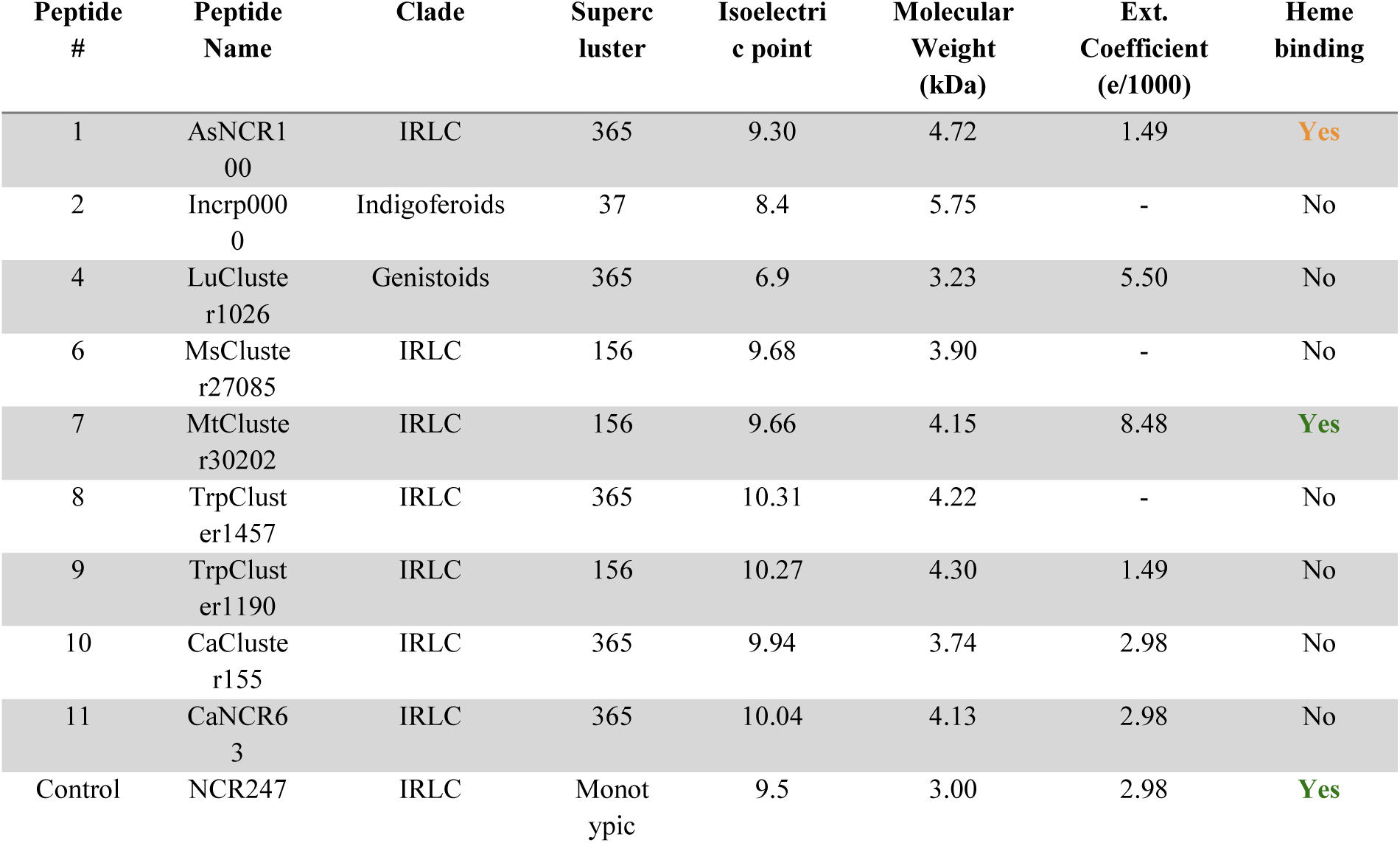
Details about the NCR peptides functionally tested in this study.

Physicochemical properties and heme binding activity of the functionally tested NCR peptides. Each peptide is identified by its assigned number (Peptide#) and name (Peptide Name). Clade indicates the phylogenetic grouping of the peptide within the legume phylogeny, and Supercluster refers to the structural cluster classification. Isoelectric point (pI) is the theoretical pH at which the peptide carries no net charge. Molecular Weight (kDa) is calculated based on the peptide’s amino acid sequence. Ext. Coefficient (e/1000) denotes the molar extinction coefficient per 1000 Da at 280 nm, reflecting aromatic amino acid content. “-” indicates that no reliable value could be determined. Heme binding specifies whether the peptide binds heme or not. The green color means it binds to heme, and the orange means it binds to a lesser extent. NCR247 is a Control reference peptide with known antimicrobial and heme-binding activity.

First, we checked the anti-bacterial activity against a gram-negative bacterium, *E. coli*, and a gram-positive bacterium, *Bacillus subtilis*. The native bacterium on which NCR247 acts during symbiosis is the rhizobial bacterium *S. meliloti*. To compare the activity to NCR247, we also tested the growth inhibition induced by these peptides on *S. meliloti*. Next, we tested at sublethal concentrations to see if these peptides could induce polyploidy in *S. meliloti* since it is an NCR-dependent characteristic feature of differentiated bacteroids. Ploidy was determined by directly measuring the DNA content of synchronized, peptide-treated cells through flow cytometry. Multivariate analysis of the three relevant antimicrobial data and ploidy data of *S. meliloti* revealed that peptides clustered by functional activity rather than structural supercluster clustering (**Figure 6**). As expected, the nine tested NCR peptides exhibited antimicrobial activity against *S. meliloti*, but to different extents. The neutral NCR peptide from *L. luteus* exhibited less antimicrobial activity (**Figure 6, S8**), while 7 of the eight cationic NCR peptides inhibited *S. meliloti* growth (**Figure 6, S8**). These seven NCR peptides were also toxic to *E. coli* (**Figure 6, S8**) but to various extents. *B. subtilis* 3610 was resistant to almost all the tested NCR peptides (**Figure S8**), while the growth of *B. subtilis* PY7 showed a similar profile to that of *E. coli*, where the cationic NCR peptides inhibited its growth (**Figure 6, S8**). At lower peptide concentrations, polyploidy was induced in *S. meliloti* by the same seven cationic peptides that displayed antibacterial activity (**Figure 6, S10**). The neutral *L. luteus* NCR peptide did not induce changes in DNA content (**Figure 6**), yet it is differentially and highly expressed in nodules; we suggest that this NCR peptide is essential in the presence of other NCR peptides. The cationic *T. pratense* NCR peptide also did not induce an increase in the DNA content of *S. meliloti* (**Figure 6**). Notably, despite clustering structurally with defensin-like peptides in the SC_37 supercluster, the *I. argentea* NCR peptide exhibited moderate antimicrobial activity across all tested bacteria and induced significant ploidy changes in *S. meliloti* (4.9-fold increase over untreated) (**Figure 6, S8, S10**). Thus, phenotypes of the newly identified NCR peptides are closely aligned with what is known for well-studied NCR peptides of *M. truncatula*. This functional convergence demonstrates that NCR peptide functions have evolved independently from diverse structural backbones, including defensin peptides.

**Figure 6.**
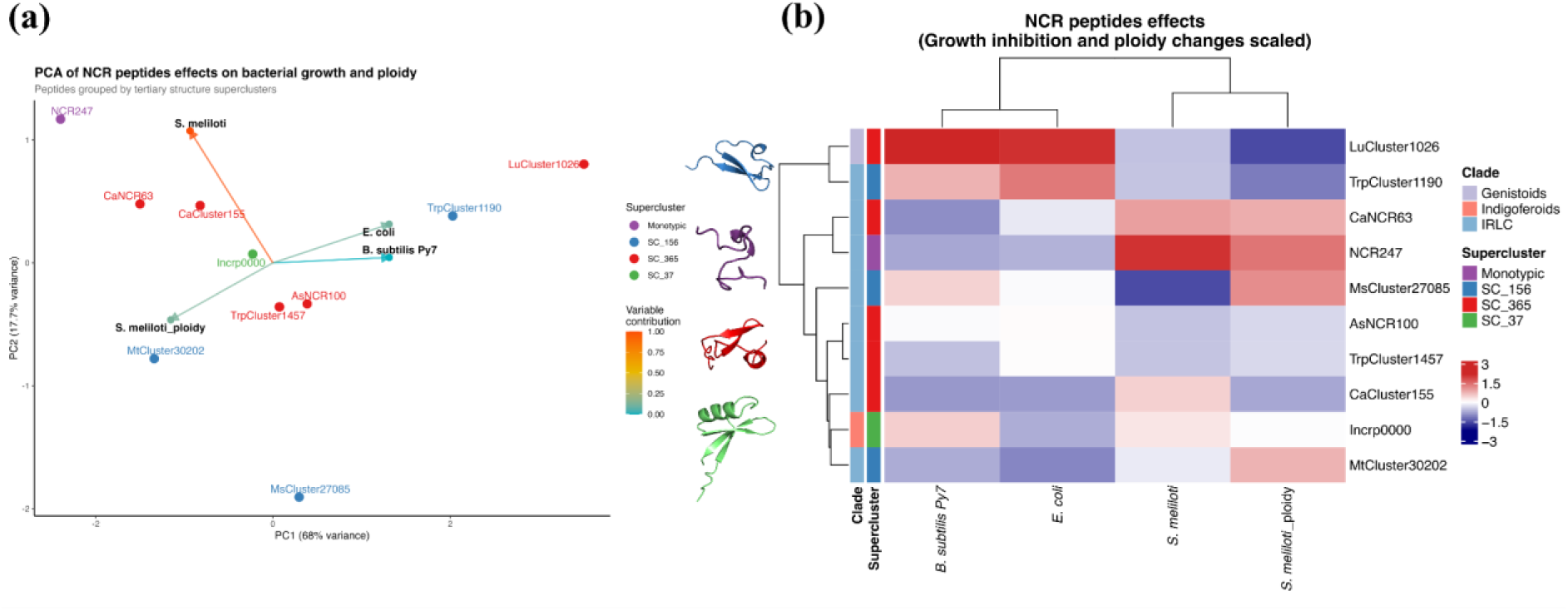
Integrated analysis of NCR peptide effects on bacterial growth and ploidy with structural supercluster classification. **(a)** Principal Component Analysis (PCA) biplot showing the distribution of NCR peptides based on their effects on bacterial growth inhibition (calculated using the mean area under curve of the three replicates and relative to the untreated bacteria) across the three bacterial strains (*B. subtilis* Py7, *E. coli*, and *S. meliloti*) and ploidy changes in *S. meliloti* (percentage of polyploid bacteria normalized to be relative to the untreated bacteria). PC1 and PC2 axes show the percentage of variance explained. Peptides are colored according to their structural supercluster membership: Monotypic (purple), SC_156 (blue), SC_365 (red), and SC_37 (green). Representative tertiary structures for each supercluster are displayed, showing the conserved cysteine-rich β-sheet core structure with variations in loop regions and terminal extensions. Variable contributions are overlaid as loading arrows originating at the origin. Arrow direction indicates the correlation of each variable with PC1/PC2, and arrow length reflects its magnitude. Arrows are colored by contribution to PC1+PC2 (from blue (low) to red (high)). **(b)** Hierarchical clustering heatmap displaying normalized bacterial responses to NCR peptide treatments. Data were independently Z-score normalized for each measurement type to enable cross-species comparison. The color gradient represents scaled effects: for growth data, blue indicates strong growth inhibition (effective antimicrobial activity), white represents the mean effect across all peptides tested, and red indicates minimal to no growth inhibition (similar to untreated controls); for ploidy data, blue indicates low ploidy changes (similar to untreated controls), white represents the mean ploidy effect, and red indicates high ploidy induction (strong endoreduplication effect). The intensity scale ranges from -3 to +3 standard deviations. Hierarchical clustering was performed using Ward’s linkage method on Euclidean distances. The dendrogram (left) shows peptide clustering patterns. Annotation bars indicate the structural supercluster and the clade of the peptides.

Our previous findings demonstrated that, within *M. truncatula*, heme binding is highly selective, with only NCR247 exhibiting this property, while other cationic peptides like NCR035 do not bind heme (Sankari et al., 2022). When tested for *in vitro* heme binding activity, one other *M. truncatula* NCR peptide bound heme with an absorption maxima at 420 nm (**Figure S9, Table 1**). This differs from NCR247, which binds heme with 360 and 450 nm absorption maxima. (**Figure S9, Table 1**). Interestingly, the *A. sinicus* NCR peptide from another supercluster also bound heme with an absorption maximum at 420 nm, but to a lesser extent (**Figure S9, Table 1**), reflecting a convergent trend in heme binding capability. This suggests functional specialization among NCR peptides, with some potentially involved in iron homeostasis during symbiosis, while others may have alternative functions.

## DISCUSSION

This study was motivated by multiple questions about the evolution and function of NCR peptides in legume-rhizobium symbiosis. First, the remarkable observation that Terminal Bacteroid Differentiation (TBD) occurs in five phylogenetically diverse legume clades (Oono et al., 2010) raised fundamental questions about whether this trait evolved once or multiple times independently (Oono et al., 2010). Second, NCR peptides have only been characterized in two of the five clades (IRLC and Dalbergioids), which leaves a significant gap in our understanding of TBD mechanisms across legumes (Czernic et al., 2015; Montiel et al., 2017). Third, the extreme sequence divergence of NCR peptides, even within a single species, suggests that traditional phylogenetic approaches may be inadequate for determining their evolutionary history (Alunni & Gourion, 2016). Finally, whether NCRs evolved from defensins (Salgado et al., 2022) or arose independently (Czernic et al., 2015) remained unresolved.

First, we identified conflicting reports about the emergence of TBD in the Indigoferoids and Millettioids clades, which created a paradox. Oono et al. (2010) reported TBD in *Tephrosia* species (Millettioid), but not in *Indigofera* species (Indigoferoids). However, our experiments revealed the opposite pattern. We observed clear evidence of TBD in *Indigofera argentea*. The bacteroids showed increased DNA content (3C) and slight cell enlargement, accompanied by the expression of 12 NCR peptides. In contrast, our experiments with *Tephrosia* species (data not shown) failed to detect TBD, despite extensive sampling across multiple growth conditions and rhizobial strains. This paradox suggests that TBD distribution may be more complex than previously recognized, where environmental or strain-specific factors may critically influence TBD expression.

The limited TBD observed in *I. argentea* (only reaching 3C compared to >16C in *Medicago*) (Mergaert et al., 2006) raised additional questions about the quantitative relationship between NCR peptide diversity and TBD extent. This relationship was initially proposed by Montiel et al. (2017), who identified a correlation between the extent of TBD and the number of NCR peptides across IRLC species. Furthermore, while *Lupinus* species from the Genistoid clade were previously documented to induce TBD (Oono et al., 2010), no prior studies had examined the presence of NCR peptides in these species or the potential correlation between their expression and the extent of TBD observed. These gaps in scientific knowledge underscore the necessity for a systematic, multi-clade analysis that integrates computational and experimental approaches to comprehensively characterize NCR peptides, extending beyond the well-studied IRLC and Dalbergioid clades.

To address these questions, we conducted the most comprehensive analysis of NCR peptides to date, identifying 3,710 NCR peptides across four legume clades that induce TBD. The successful application of the SPADA pipeline (Zhou et al., 2013), combined with custom HMM profiles and rigorous filtering criteria, enabled the discovery of NCR peptides in previously uncharacterized species. The identification of 12 NCR peptides in *I. argentea* provides a molecular explanation for the observed TBD in this species, while the discovery of 192 NCR peptides across three *Lupinus* species extends our understanding to the Genistoid clade. This represents a substantial advance over previous studies limited to IRLC and Dalbergioid clades (Czernic et al., 2015; Montiel et al., 2017).

The recovery rate of known NCR peptides (87-100% in well-studied species) validates our computational approach, while the discovery of 13-70% novel NCR peptides even in extensively studied IRLC species demonstrates the sensitivity of our expanded detection method. Moreover, the marked variation in NCR peptide numbers across species (from 12 in *I. argentea* to over 700 in *Medicago truncatula*) strongly correlates with the extent of TBD observed. This quantitative relationship supports the hypothesis that NCR peptide diversity drives the extent of symbiont manipulation (Van de Velde et al., 2010). Within the Genistoid clade, *Lupinus albus* harbors significantly fewer and less highly expressed NCR peptides compared to *L. luteus* and *L. mariae-josephae* (p < 0.001, Wilcoxon test), suggesting potential differences in symbiotic strategies even within a single genus.

The sequence-based clustering analysis revealed a complex pattern that challenges the simple models of NCR peptide evolution. Initial orthology analysis using OrthAgogue (Ekseth et al., 2014) and MCL clustering (Enright et al., 2002) with six species containing known NCR repertoires demonstrated complete clade-specificity, with all 651 IRLC and 20 Dalbergioid clusters showing no inter-clade members. This preliminary finding indicated the potential for distinct origins of NCR peptides across different clades. However, this perspective shifted considerably when our analysis was expanded to include Genistoid and Indigoferoid species.

The finding that 86% (75/87) of *L. luteus* NCR peptides and 88% (61/69) of *L. mariae-josephae* NCR peptides matched IRLC HMM profiles reveals unexpected sequence conservation between these phylogenetically distant clades, which diverged approximately 50 million years ago (Lavin et al., 2005). This inter-clade clustering does not necessarily indicate a single common origin but instead suggests convergent sequence evolution under similar functional constraints.

Interestingly, the widely used NCR peptide NCR247, encoded by *M. truncatula*, has homologs only in *M. sativ*a. Recently, a study that identified the NCR peptides from the transcriptome of *M. sativa* and *M. officinalis* (Huang et al., 2022) using SPADA reported that no homolog of the NCR peptides known to be essential for symbiosis in *M. truncatula* (i.e., NCR211 and NCR169) were found in *M. sativa* and *M. officinalis*. However, with our analysis, among the newly identified 633 *M. sativa* NCR peptides, we found the homologs of NCR211 and NCR169. Moreover, we also recovered in *M. sativa* and *M. officinalis* homologues of the recently identified essential NCR peptides in *M. truncatula* (Horváth et al., 2023) (NCR343 and NCR-new35). However, the lack of homologs for essential *Medicago* NCR peptides in other species reveals the paradox of NCR peptide evolution: peptides crucial for symbiosis in one species may be absent in closely related species with similar symbiotic phenotypes. This suggests massive functional redundancy within NCR peptide repertoires, where different combinations of peptides can achieve comparable outcomes. Such redundancy would provide evolutionary flexibility, allowing rapid adaptation to new rhizobial strains while maintaining symbiotic function.

Structure-based analysis has revolutionized our understanding of NCR peptide evolution, revealing relationships hidden at the sequence level. Despite an average sequence identity of only 27% within structural superclusters, the corresponding structures exhibited significant conservation, with a mean TM score of 0.75, well above the 0.5 threshold for structural similarity. Despite sequence divergence, this substantial conservation of tertiary structure explains why traditional, sequence-based phylogenetic approaches failed to capture the evolutionary history of NCR peptides.

The Foldseek clustering regrouped our 396 NCR sequence-based NCR clusters and 48 defensin clusters into 23 structural superclusters, with 14 containing members from multiple clades. This structural convergence suggests that NCR peptides across different legume lineages have evolved to adopt similar three-dimensional conformations, likely reflecting shared functional constraints in bacteroid manipulation. Structure-based clustering and phylogeny reveal two primary evolutionary paths. Dalbergioid and Indigoferoid NCR peptides derive from defensin ancestors, maintaining the characteristic eight-cysteine pattern and defensin fold. In contrast, IRLC and Genistoid NCRs form a structurally distinct group with no clear relationship to known protein families. This bifurcation cannot be explained by divergence from a common ancestor but instead indicates independent recruitment of different protein scaffolds for similar functions.

We provide direct evidence for convergent evolution through our functional validation of nine NCR peptides from different clades and structural superclusters. Multivariate analysis revealed that functional properties cluster independently of structural superclusters. Peptides from different superclusters showed similar activity profiles. This functional convergence across structural classes suggests that legumes have evolved multiple molecular solutions to controlling their bacterial symbionts independently. Despite its high nodule expression, the lack of antimicrobial activity of the neutral *L. luteus* peptide suggests that some NCR peptides may function primarily in combination with others or may have non-antimicrobial roles in symbiosis.

By synthesizing our sequence, structural, and functional data, we support convergent evolution through at least two independent origins of NCR peptides. Dalbergioid and Indigoferoid NCR peptides evolved from defensins, maintaining the 8-cysteine pattern and defensin fold while gaining symbiotic functions, as supported by their structural clustering with defensins and demonstrated ability to induce TBD features. Conversely, IRLC and Genistoid NCRs represent independent evolution from an unknown ancestor or de novo origin, shown by their distinct structural clustering, 4- and 6-cysteine motifs, and lack of similarity to known proteins. However, this pattern contradicts the actual legume phylogeny, where Dalbergioids group with Genistoids, and IRLC with Indigoferoids (Lavin et al., 2005; Oono et al., 2010). This phylogenetic incongruence suggests multiple independent recruitments of similar peptide types across the legume tree, rather than simple inheritance from a common ancestor.

The rapid sequence evolution observed within structural superclusters suggests that NCR peptides experience relaxed selective constraints on primary sequence if structural integrity is maintained. This evolutionary flexibility may allow fast adaptation to new bacterial partners or environmental conditions while preserving core functional capabilities. The presence of species-specific NCR peptide clusters alongside conserved inter-clade clusters indicates ongoing birth-and-death evolution, with new peptides constantly arising through duplication and divergence while others are lost or pseudogenized.

Finally, through this study, we demonstrated the importance of integrating sequence-based, statistical, and structural analysis and how this method can reveal novel evolutionary insights about NCR peptides hidden by their sequence divergence. The successful application of AlphaFold2 predictions combined with Foldseek clustering and Foldtree phylogenetics to analyze thousands of small peptides demonstrates the scalability and reliability of this approach. This methodology could be applied to other families of small, rapidly evolving peptides, including antimicrobial peptides, toxins, hormones, and signaling molecules across diverse biological systems. Moreover, our findings have significant implications for agricultural biotechnology and sustainable agriculture. Understanding the molecular basis of enhanced nitrogen fixation through TBD could allow us to engineer more efficient symbioses in crop legumes that lack NCR peptides.

## METHODS

### Bacterial strains, nodulation assays, and analysis

*Sinorhizobium meliloti* 1021 strain was grown at 28°C in YEB (0.5% beef extract, 0.1% yeast extract, 0.5% peptone, 0.5% sucrose, 0.04% MgSO_4_ 7H_2_O, pH 7.5) or LBMC (LB medium supplemented with 2.5 mM CaCl_2_ and 2.5 mM MgSO_4_). *Bradyrhizobium elkanii* SA281 strain was cultivated at 28°C in YM (Vincent, 1970) medium. Streptomycin (Sm; 500 µg/mL) was added to the media as appropriate.

Seeds of *Indigofera argentea,* from an accession collected in 2010 in the Jizan desert in Saudi Arabia, which is part of the Nagoya protocol since 2020, were provided by Ton Bisseling. Seeds were treated with 96% sulfuric acid for seven minutes before being rinsed six times with double-distilled water. The seeds were then surface sterilized with 4% commercial bleach for 10 minutes and rinsed seven times before being soaked in sterile double-distilled water for three hours at room temperature in the dark. The sterilized seeds were plated on water agar in 9 cm plates and incubated at 4°C for 12 hours in the dark and then at 28°C for 24 hours in the dark. After that, the seeds were exposed to light for 4-5 days, and then the germinated seeds were planted in perlite/sand (2:1 vol/vol), humidified with nutrient water in 1.5L pots in the greenhouse (28°C, 16 hours of light and 8 hours of dark, humidity 60%). The seedlings were grown in the greenhouse for 3-4 days without watering and then inoculated with 20 mL per pot of *B. elkanii* (SA281) at an OD_600nm_ of 0.05 and grown for another 2-3 days without watering. The plants were watered every three days, alternating tap water and a commercial N-free fertilizer (PlantProd solution [N-P-K, 0-15-40; Fertil] at 1 g per liter) (Kazmierczak et al., 2017). Root nodules from inoculated plants and roots from uninoculated plants were collected 8 weeks post-inoculation, immediately frozen with liquid N_2_, and stored at -80°C until use. In addition, fresh nodules (non-frozen) were kept for confocal microscopy and flow cytometry experiments.

### RNA extraction and sequencing

The total RNA from three biological replicates of frozen root and nodule tissue of *Indigofera argentea* was extracted using the MasterPure Complete DNA and RNA purification kit (Epicentre, Madison, WI, USA), following the manufacturer’s protocol. DNA was removed using a Turbo DNA-free kit from Ambion (Ambion, Thermo Fisher Scientific, Waltham, MA, USA), treating one µg of RNA per reaction to degrade any contaminating DNA. The concentrations of the purified RNA samples were measured using a DeNovix Spectrophotometer DS-11. RNA integrity was assessed with gel electrophoresis.

Library preparation and Illumina sequencing were performed at “Plateforme de Séquençage Haut Débit I2BC” (Gif-sur-Yvette, France). Before library preparation, the quality of the RNA samples was assessed with an Agilent Bioanalyzer RNA 6000 pico chip, and the RNA concentrations were measured with a Qubit fluorometer. RNA library preparation was performed using Illumina Stranded mRNA Prep with ribosomal RNA depletion and PolyA purification. The libraries were sequenced on Illumina NextSeq 2000 to generate paired-end reads of 150x2 bases. The raw data were demultiplexed using bcl-convert 4.1.5, and then the adapters were trimmed using cutadapt 3.2 (Martin, 2011).

### Transcriptome *de novo* assembly and annotation

We preprocessed the newly obtained raw reads (root and nodule *I. argentea*) and publicly available SRR datasets (nodules of *Lupinus mariae-josephae* and *L. luteus*) to ensure high-quality data for the downstream analysis. Briefly, the remaining adaptors were removed with Fastp version 0.20.0 (Chen, 2023), reads with unfixable errors removed with Rcorrector version 1.0.4 (Song & Florea, 2015) and the FilterUncorrectablePEfastq.py python script (github.com/harvardinformatics/ TranscriptomeAssemblyTools/), and the remaining short or low-quality reads (Q score < 20 and length < 25) were removed with TrimGalore version 0.6.6. We assessed the read quality after each preprocessing step with FastQC (Andrews, 2010). To remove possible bacterial contamination from *I. argentea* root and nodule read sets, we mapped our reads using Bowtie2 version 2.3.5.1 (Langmead & Salzberg, 2012) against the *Bradyrhizobium elkanii* SA281 bacterial strain we used to inoculate our plants. We then used Samtools version 1.10 (Danecek et al., 2021) to keep only unmapped reads and convert our data to Fastq format.

The transcriptomes of all four RNA-seq datasets were de novo assembled separately following the same process. First, we simultaneously provided our processed clean reads of the three replicates to Trinity version 2.6.6 (Grabherr et al., 2011) to generate a de novo transcriptome assembly. Second, to merge the *Indigofera argentea* roots and nodules contigs, we used CD-HIT version 4.8.1 (Fu et al., 2012). Then, we clustered our contigs into gene-level clusters, first using Corset version 1.0.9 (Davidson & Oshlack, 2014) to find gene isoforms and then Lace version 1.14.1 (Davidson et al., 2017) to merge those isoforms to form supertranscripts. The nodule transcripts of the two Lupinus species were also clustered into supertranscripts using the same approach (Corset and Lace).

We checked the quality of the Trinity and SuperTranscript assemblies using STAR version 2.7.3a (Dobin et al., 2013) to obtain the alignment rate and BUSCO version 5.5.0 (Simão et al., 2015) to assess the completeness of the assemblies using ‘Viridiplantae’ and ‘Fabales’ databases. For *I. argentea*, we checked the quality of the assemblies (alignment rate and completeness) before and after merging the root and nodule assemblies to check the quality of the merged assembly.

We predicted coding regions using TransDecoder version 5.7.1 (*TransDecoder/TransDecoder: TransDecoder Source*, n.d.)based on the results of blastp version 2.12.0+ (Camacho et al., 2009) searches against the UniProt database (The UniProt Consortium, 2019)(release-2023_05) supplemented with all the known NCR peptides (Czernic et al., 2015; Montiel et al., 2017; Raul et al., 2021). Finally, we performed a functional annotation of the *I. argentea* protein sequences using the GFAP (Gene Functional Annotation for Plants) pipeline using *Glycine max* (the closest reference plant species available in the database) (Xu et al., 2023). The AHL and LegHB proteins were annotated manually using tblastn and all AHL and LegHB from legumes from NCBI because these genes function as positive controls, as leghemoglobin proteins (LegHB) are crucial for symbiotic nitrogen fixation and are induced explicitly in nodule development (Ott et al., 2005). At the same time, AHL transcription factors are essential for NCR gene expression and nodule development in legumes that form nitrogen-fixing partnerships with rhizobia (S. Zhang et al., 2023).

### Homology, orthology, and clustering analysis

The genomes and predicted proteomes of legume species with known NCR peptides (four IRLC and two Dalbergioid species) were collected from NCBI and the Legume Information System (LIS, https://www.legumeinfo.org/) (Berendzen et al., 2021). The known IRLC and Dalbergioid NCR peptide sequences were collected from (Montiel et al., 2017) and (Czernic et al., 2015; Raul et al., 2021), respectively.

To ensure the presence of NCR peptides in the predicted proteomes, we searched for the known NCR peptides in their corresponding species proteome using blastp (Camacho et al., 2009). If the NCR peptides were not found in the proteome, tblastn was used to search for them in their corresponding genome. If found in the genome, the NCR peptides were added to the species’ proteome.

Using the species proteomes, including all known NCR peptides, we scored the similarity among all species sequences using blastp (Camacho et al., 2009). We used those scores to define a set of orthologs using orthAgogue software (Ekseth et al., 2014) and then regrouped those orthologs into clusters using Markov Clustering (MCL) (Van Dongen, 2008). Custom scripts, available through GitHub (https://github.com/amira-boukh/Legume_Transcriptome_Assembly_and_NCR_identification), were used to extract NCR clusters from all protein clusters. Any cluster containing at least two NCR peptides was defined as an “NCR cluster”, while clusters with at least one NCR peptide and at least one additional protein were described as “NCR-mixed clusters”. We extracted only the NCR peptides for downstream analyses in the NCR-mixed clusters with at least two NCR peptides.

### NCR peptide detection and classification

The DNA and protein sequences of NCR clusters were extracted using custom scripts. SignalP version 4.0 (Petersen et al., 2011) excluded clusters with no signal peptide in at least two NCR peptides. Macse2 produced codon-based multiple sequence alignments from CDS sequences of the remaining NCR clusters. Translated protein Hidden Markov Models (HMM) profiles were built from those alignments with hmmbuild from HMMER version 3.3.2 (Eddy, 1998).

The legume species used to search NCR peptides were the above *de novo* assembled transcriptomes (*I. argentea*, *L. luteus*, and *L. mariae-josephae*), the assembled nodule transcriptomes of *M. sativa* (Huang et al., 2022), *M. officinalis* (Huang et al., 2022), and *C. arietimum* (Kant et al., 2016), and the publicly available legume genomes and nodule RNA-sequencing data for *Medicago truncatula*, *Pisum sativum*, *Trifolium pratense*, *Arachis hypogaea, Aeschynomene evenia, Lupinus albus, Lotus japonicus, Cajanus cajan, Phaseolus vulgaris, Vigna angularis,* and *Glycine max* (Alves-Carvalho et al., 2015; Clevenger et al., 2016; Keller et al., 2018; O’Rourke et al., 2014; Pazhamala et al., 2017; Pecrix et al., 2018; Quilbé et al., 2021; Sakai et al., 2015; Trujillo et al., 2019; Yuan et al., 2021).

The RNA-seq data were downloaded from the Sequence Read Archive (SRA) database. They were converted to fastq files using the fastq-dump tool from the SRA Toolkit. Their corresponding genomes were also downloaded from NCBI and LIS. The accession numbers of the genomes and RNA-seq data are listed in **Dataset S1**.

As described above, NCR peptides are small and highly divergent sequences. Therefore, known functional annotation methods are not sensitive enough to detect NCR peptides from genomes or transcriptomes. Consequently, we used SPADA (Small Peptide Alignment Discovery Application) pipeline version 1.0 (Zhou et al., 2013) to search for NCR peptides in our genomes and nodules transcriptomes. SPADA is a computational pipeline that, when provided with multiple sequence alignments for a particular gene family, identifies all members of this family in a target genome sequence. The SPADA pipeline specializes in predicting cysteine-rich peptides in plant genomes. First, we used SPADA’s “seq.check” command to check our genomes. Second, we used the command “build_profile” to build HMM profiles from our IRLC and Dalbergioid NCR cluster alignments. We then ran SPADA three times separately for each genome or assembled transcriptome, one using IRLC profiles, one using Dalbergioid profiles, and one using CRP profiles from the SPADA pipeline, which are plant cysteine-rich peptides that we used because we were not sure that our clade-specific clusters could capture all NCRs in other clades. For each genome, we merged the results from the three analyses, converted the merged results to fasta format, and removed the duplicates.

The predicted putative NCR peptides were filtered according to their length, cysteine motif, and nodule expression. To predict the length of mature peptides, we used SignalP version 4.1g (Petersen et al., 2011) with the “notm” network to predict the cleavage sites and extract the mature peptides. Using custom scripts, the length of the mature peptides and the number of cysteines were counted, and we kept only NCR peptides whose mature peptides have fewer than 100 aa and at least four cysteines. We then assessed the nodule expression by mapping the nodule RNA-seq data’s reads against the genome using STAR version 2.7.3a (Dobin et al., 2013), and quantifying the number of reads using htseq-count version 0.12.3 (Putri et al., 2022). We kept only putative NCR peptides expressed in nodules (See Differential expression analysis).

We classified the retained NCR peptides into two groups: NCR peptides with four or six cysteines (NCR-motif) and NCRs with eight cysteines (defensin-motif). For the IRLC species, we annotated as NCR peptides only those with the NCR motif, as defensin motif-containing peptides are presumed to act in the innate immune system of IRLC plants. However, we annotated NCR-motif and defensin-motif as NCR peptides for the other clades. To compute the pI values of the NCR peptides and clusters, we used the R package pIR, where the pI of each peptide was calculated based on the mean values from all prediction methods, excluding the highest and lowest values (Audain et al., 2016).

We then performed a classification step for NCR peptides found with CRP (Cysteine-Rich Peptide) profiles (SPADA did not classify them with our IRLC or Dalbergioid profiles), where we searched them against our cluster profiles using hmmsearch version 3.3.2 (Johnson et al., 2010) and chose the best hit profile to classify each sequence. Finally, we merged NCR peptides classified with SPADA and those classified with the HMMsearch approach for each species.

Moreover, for *I. argentea,* where only a few NCR peptides were found, and all of them were unclassified, we ran SPADA a second time using our expanded clusters, including newly found NCR peptides in IRLC, Dalbergioids, and Genistoids, and also using one NCR HMM profile of *I. argentea* that we built using the five NCR peptides found in (Ren, 2018). The filtration steps here were more flexible, where we did not exclude long NCR peptides that are differentially expressed in nodules and have an NCR cysteine motif.

### Differential expression analysis

To estimate the gene expression levels of *I. argentea* and to highlight the differentially expressed NCR peptides between root and nodules, we used the mapping-based mode of salmon version 0.12.0 (Patro et al., 2017) to map our samples individually to the reference transcriptome described above. We used the R package DESEQ2 version 1.32.0 (Love et al., 2014) to predict the differentially expressed genes between root and nodules of *I. argentea* using the raw counts for each replicate found with salmon, the length of each gene for normalization, and the annotation of the genes. We also used the diCoexpress R package (Lambert et al., 2020), which performs differential expression analysis using generalized linear models with the edgeR package (Robinson et al., 2010). The genes with log2(FoldChange) > 1 and the adjusted p-value < 0.05 were considered up-regulated, and the genes with log2(FoldChange) < -1 and the adjusted p-value < 0.05 were considered down-regulated.

For all other legume species where SPADA was run on the genomes or nodules transcriptomes, we performed a differential expression analysis to assess the expression of NCR peptides in the nodules. First, we used STAR version 2.7.3a (Dobin et al., 2013) in two-pass mode to align the roots and nodule reads (fastq files) on the genome (nodule transcriptome). In the first pass, splice junctions were identified to guide a second alignment, enhancing mapping accuracy. Then, the final alignments from the second pass were used for transcripts quantification using htseq-count version 0.12.3 (Putri et al., 2022). Using custom scripts, we extracted NCR transcripts that were differentially expressed on nodules and used them to filter out the other putative NCR peptides.

### Structural and phylogenetic analysis

A complete command-line local installation of Alphafold2 version 2.2 (Jumper et al., 2021) on the I2BC server was used to predict the tertiary structures of one randomly picked NCR peptide per cluster. Structural models were constructed using Alphafold2’s own embeddings, supplemented with macse-generated multiple sequence alignments of sequence-based clusters. All templates downloaded on 30 June 2022 were allowed for structure modeling. Five models were predicted, and we selected the model with the best pLDDT (predicted Local Distance Difference Test) score. We kept only the NCR structures with a pLDDT score higher than 70 for further analysis. For the non-classified predicted NCR peptides from Indigoferoids (*Indigofera argentea)* and Genistoids (*Lupinus* spp.), we first used CD-HIT version 4.8.1 (Fu et al., 2012) with an identity threshold of 90, 80, 70, and 50% to regroup them into clusters. However, all the non-classified NCR peptides were monotypic, sharing no similarities. Thus, we predicted the tertiary structure of each NCR peptide using AlphaFold2, as described above, and we included them for further analysis in our dataset of the NCR clusters as monotypic NCR peptides. To check if NCR peptides evolved from defensins, we predicted the tertiary structures of one defensin from each of the 48 defensin orthologous clusters, ensuring to include at least one representative from each legume species, and we included them in our structural comparison analysis.

We then used Foldseek version 4-645b789 (van Kempen et al., 2024), a structure clustering approach, to regroup all the high-quality NCR and defensin structures into superclusters. Foldseek is a fast method that converts the tertiary structures into 1D vectors that contain the structure information, which decreases computation times by four to five orders of magnitude. The TM scores within each supercluster and between each supercluster and all others were also computed with Foldseek to compare intra-supercluster versus inter-supercluster TM scores, and the distributions of TM scores were plotted in R with ggplot. For each supercluster, all structures’ structural alignment was computed using USalign (C. Zhang et al., 2022) or MUSTANG (Konagurthu et al., 2006). These structural alignments were represented with a sausage representation with a blue-red color gradient that represents the TM score ranging from 50 to 100 of the all-vs-all alignment of all the structures of the superclusters, but defined by the alignment of all structures to the longest structure (the TM scores are the scores from all-vs-all alignment, but for a better visualization we kept only the alignment of all structures to the longest structure).

Then, to generate a structural “phylogenetic” tree based on the structural distances of all the NCR peptide and defensin structures, we used Foldtree (Moi et al., 2023) for each supercluster. Foldtree uses the all-vs-all comparison of Foldseek and creates a distance matrix from the Fident scores (sequence similarity after aligning with the structural alphabet) of Foldseek output, which is used as the input of quicktree (Howe et al., 2002) to generate the structure-based tree. The Foldseek parameters used for Foldtree were 0.5 of coverage, 0.25 of Foldseek alphabet sequence identity, an e-value of 0.1, and an exhaustive search. These parameters were chosen based on the separation of the superclusters and the congruence of the tree after testing more than 100 combinations of parameters. Furthermore, we excluded the monotypic superclusters to avoid any noise in our analysis and to have congruent trees. The trees were annotated and visualized with iTol version 5 (Letunic & Bork, 2021).

A sequence similarity network of the representative sequence of each cluster (the same predicted by Alphafold2 used for the structural analysis) was also produced using CLANS (Cluster Analysis of Sequences) (Frickey & Lupas, 2004). The network was annotated manually with the superclusters, and the same annotation was used in the structural phylogenetic tree.

### Flow cytometry

Bacteroid extraction was performed as described before (Mergaert et al., 2006). Bacteroids and free-living *B. elkanii* SA281 bacteria were stained with 5 μg/ml DAPI in BEB. After 10 minutes of incubation at room temperature, the bacteria and bacteroids were processed with a Cytoflex S cytometer (Beckman-Coulter). The data analysis was performed using cytExpert software v2.5 and the flowCore (Ellis et al., 2019) package in R.

### Confocal microscopy

Nodule live-dead imaging was performed on an SP8X confocal DMI 6000 CS inverted microscope (Leica). First, fresh nodules were harvested, embedded in 6% agarose, and sliced into 70µm slices using a Leica vibratome. The slices were incubated for 15 minutes in a 50mM Tris-HCl buffer with 0.01% calcofluor white (to stain plant cells), M2R (Sigma), 0.5 μl of Syto9 (to stain live bacteria), and 0.5 μl of Propidium iodide (to stain dead bacteria). Then, the washed sections were observed with ×10 dry and ×63 oil immersion objectives. The analysis of images was performed using ImageJ software (Collins, 2007).

### *In vitro* NCR sensitivity assays

All chemically synthesized peptides were purchased from Genscript. The purity of all peptides was > 98%. The heme assays were performed as described in (Sankari et al., 2022). The growth assays of *S. meliloti* and *E. coli* were also described in (Sankari et al., 2022). Briefly, overnight cultures were washed and diluted in GSY media. The diluted cultures were distributed in sterile 96-well plates, and optical density was measured at 600 nm every hour using a Tecan SPARK 10M microplate reader with continuous shaking at 150 rpm. To check the effect of NCR peptides on the cell cycle (i.e., genome amplification), we quantified the DNA content of *S. meliloti* supplemented with each NCR peptide using flow cytometry as described previously (Haag et al., 2011). Principal Component Analysis (PCA) and hierarchical clustering were performed on integrated data comprising bacterial growth inhibition (measured as area under the curve relative to untreated controls) for *B. subtilis* Py7, *E. coli*, and *S. meliloti*, along with ploidy changes in *S. meliloti* (percentage of >3C cells at 210 minutes with four µM peptide treatment, expressed as fold-change over untreated), and PCA was performed using the FactoMineR package (Lê et al., 2008) (v2.4) in R (v4.2.0). For the heatmap, all variables were Z-score normalized (z = (x - μ)/σ) to account for different measurement scales. Hierarchical clustering employed Ward’s minimum variance method (ward.D2) with Euclidean distances on the scaled data matrix, with visualization generated using pheatmap (Kolde, 2010) (v1.0.12) with a diverging blue-white-red color scheme representing -3 to +3 standard deviations.

## Data availability

All publicly available genome and transcriptome sequences and assembled transcriptome accessions and publications used in this study are provided in the Dataset S1. The raw sequencing data generated here are available in the Short Read Archive (SRR30017674, SRR30017696, SRR30018070, SRR30035742, SRR30035741, and SRR30035750) under the BioProject PRJNA1140874 at NCBI. The de novo transcriptome assemblies are deposited in the Transcriptome Shotgun Assembly (TSA) Sequence Database (GKXU00000000 for *I. argentea* roots assembly, GKXT00000000 for *I. argentea* nodules assembly, and GKXV00000000 for roots and nodules assembly) under the BioProject PRJNA1140874 at NCBI. The assembly stats and the GO annotation of *I. argentea* are provided in the Dataset S2. The number of NCR peptides per clade and species, their length, their NCR cluster, and superclusters are provided in Dataset S3. The amino acid sequences of all NCR peptides used and predicted in this study, classified by cluster and supercluster, are provided in Dataset S4. The Newick files of the phylogenies are available on GitHub (). All the code to replicate the analysis presented here is available through GitHub (https://github.com/amira-boukh/Legume_Transcriptome_Assembly_and_NCR_identification).

## Authors’ contributions

BA and RCRV designed the study. AB performed all bioinformatic analyses. AB, BA, and TT performed plant inoculation and analysis. BA produced the RNA-seq libraries. GdC provided bioinformatic tools. SS performed *in vitro* characterization. JS and PM helped in the interpretation. BA, JS, and RCRV supervised the study. AB wrote the first draft with extensive JS, BA, and RCRV input. All authors read and commented on the manuscript.

## ACKNOWLEDGEMENTS

A.B. benefited from a Ph.D. contract in the frame of the CNRS 80|PRIME – 2021 program (awarded to P.M.) and was partially supported by a Mitacs Globalink Research Award. B.A. benefited from a French State grant (Saclay Plant Sciences, reference n° ANR-17-EUR-0007, EUR SPS-GSR) under a France 2030 program (reference n° ANR-11-IDEX-0003). We thank Rui Huang for sharing R scripts (https://github.com/hyhy8181994/Nodule_transcriptome_script) that were used with slight modifications. The present work has benefited from Imagerie-Gif core facility supported by I’Agence Nationale de la Recherche (FBI ANR-24-INBS-0005 (BIOGEN); SPS ANR-17-EUR-0007, EUR SPS-GSR). The Stowers Institute for Medical Research funds S.S.

## Supplementary figures

**Figure S1.**
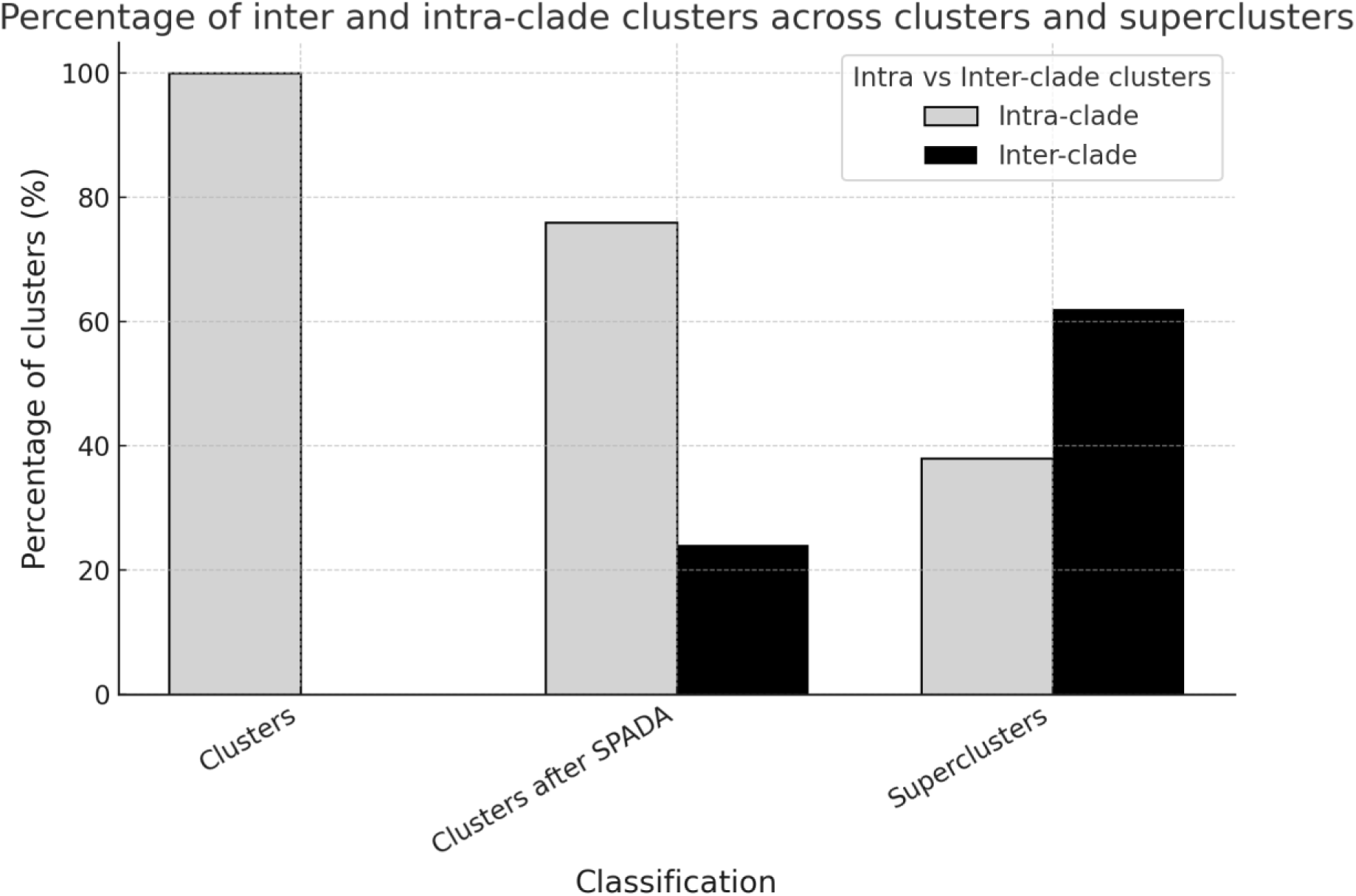
NCR clusters are clade-specific, considering IRLC and Dalbergioids, but inter-clade, considering four TBD-inducing clades after an exhaustive search with SPADA. NCR peptides have relatively conserved structures despite their sequence divergence, with more than 60% of inter-clade superclusters. The bar plots represent the percentage of clade-specific clusters versus inter-clade clusters in the first computed clusters from IRLC and Dalbergioids (left), in the reconstructed clusters from the recovered NCR peptides with SPADA (middle), and in the superclusters constructed based on 3D structures (right).

**Figure S2.**
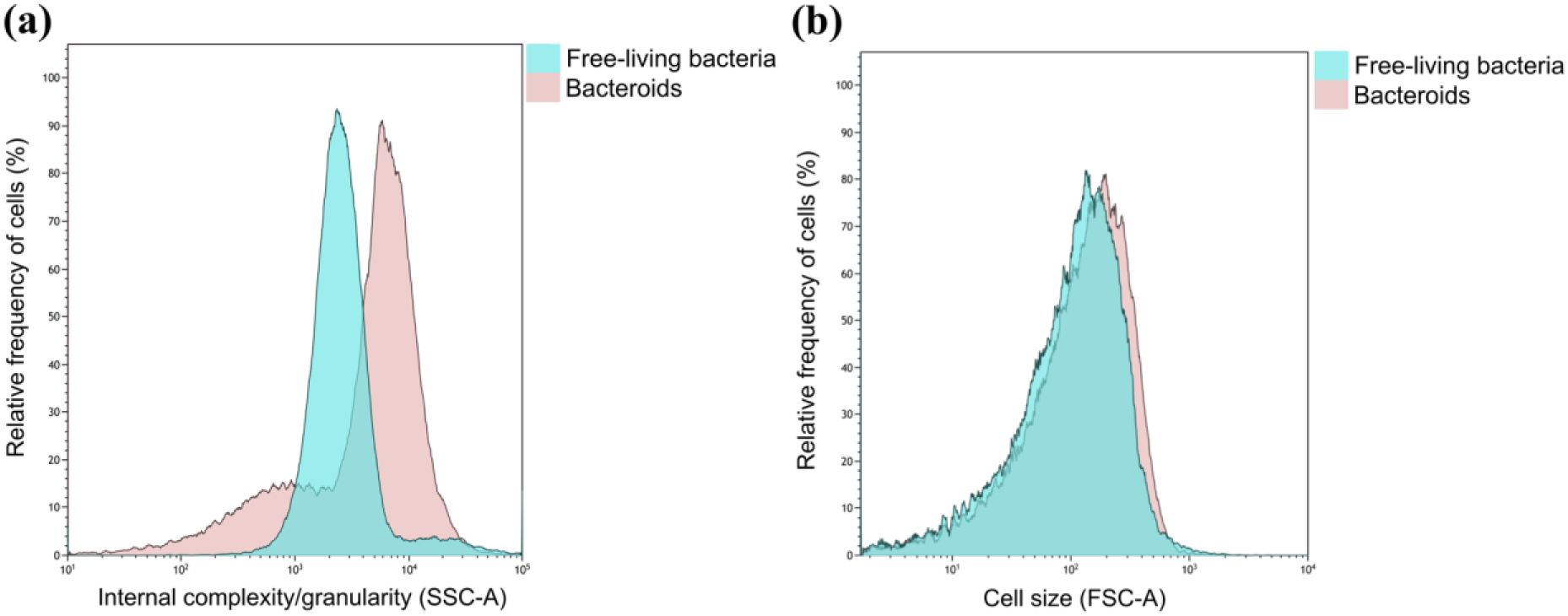
The slightly enlarged *I. argentea* bacteroids. **(a)** The granularity (estimated by SSC) of DAPI-stained *B. elkanii* strain SA281 bacteria and bacteroids isolated from *I. argentea* nodules was measured by flow cytometry. **(b)** The size (estimated by FSC) of DAPI-stained *B. elkanii* strain SA281 bacteria and bacteroids isolated from *I. argentea* nodules was measured by flow cytometry.

**Figure S3.**
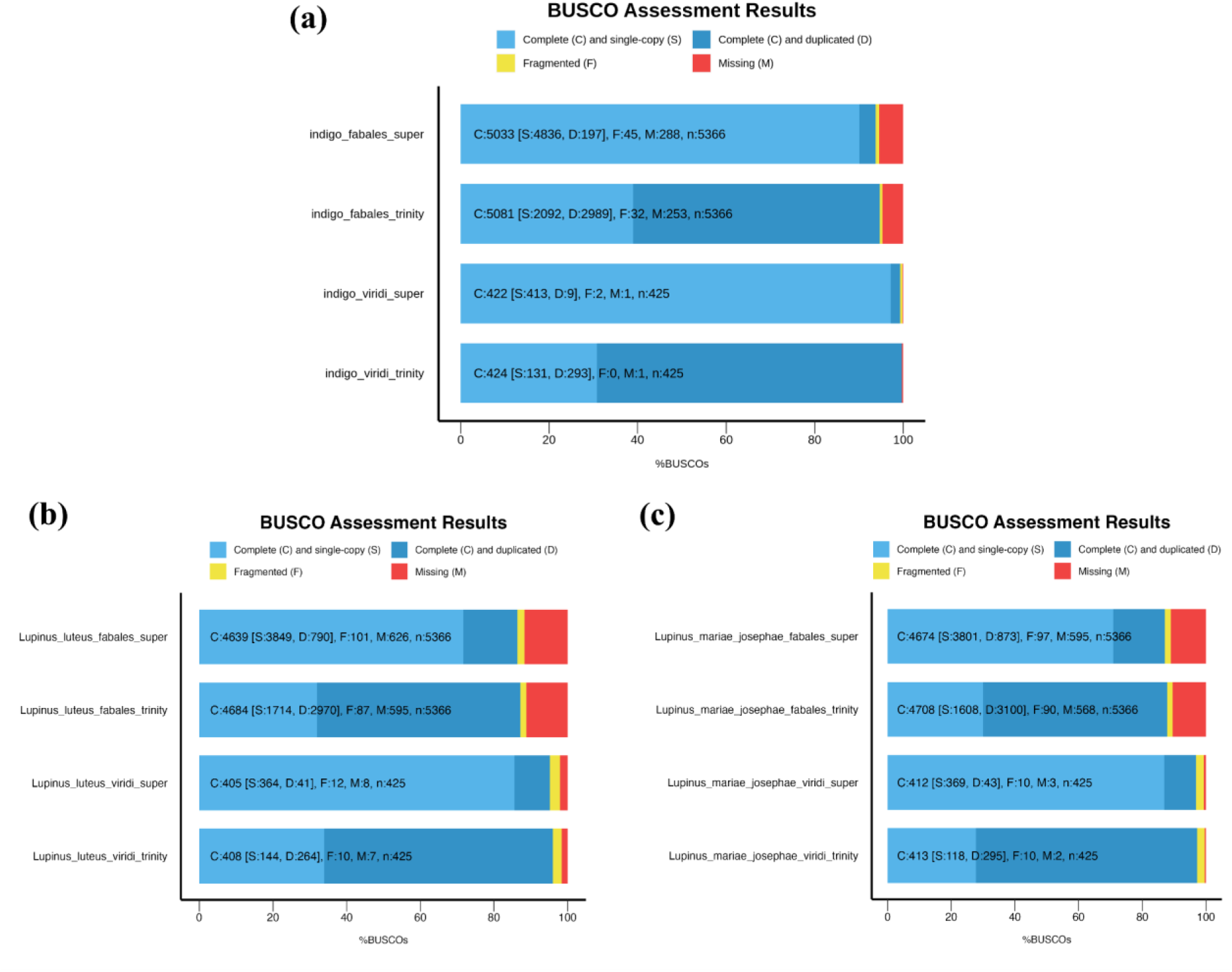
BUSCO assessment of the completeness of the *de novo* transcriptome and the supertranscripts. **(a)** In *I. argentea,* 95% of the Viridiplantae and 86% of the Fabales BUSCO genes were identified as complete and single-copy. **(b)** In *L. luteus,* 86% of the Viridiplantae and 71% of the Fabales BUSCO genes were identified as complete and single-copy. **(c)** In *L. mariae-josephae,* 87% of the Viridiplantae and 71% of the Fabales BUSCO genes were identified as complete and single-copy.

**Figure S4.**
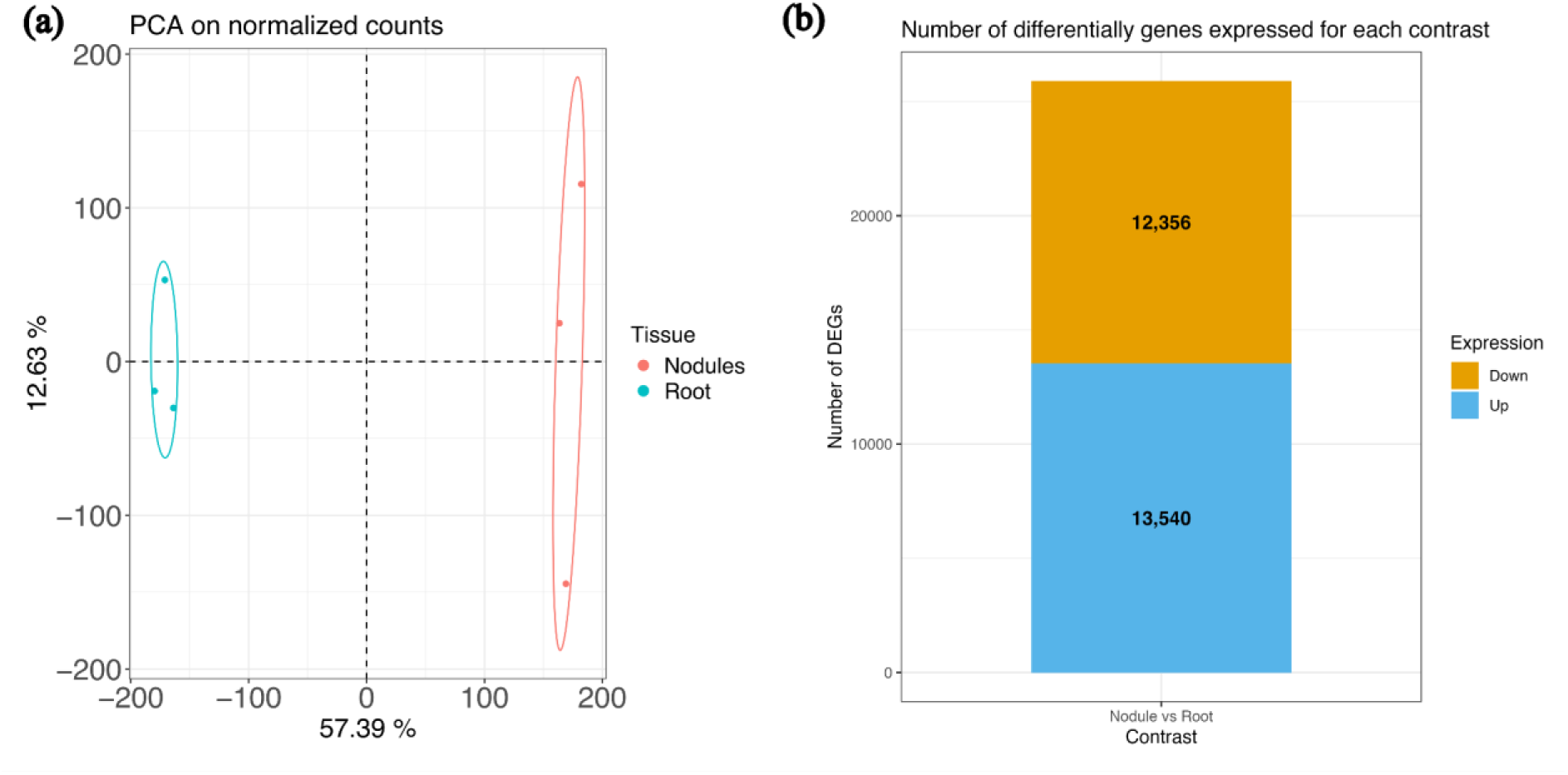
Quality control of the RNA-seq dataset and the number of up- and down-regulated genes in nodules compared to roots in *I. argentea.* (a) Principal Component Analysis (PCA) on the normalized counts of *I. argentea* to assess the quality of the roots and nodules RNA-seq datasets. (b) The number of up- and down-regulated genes in nodules of *I. argentea* compared to the roots (non-inoculated).

**Figure S5.**
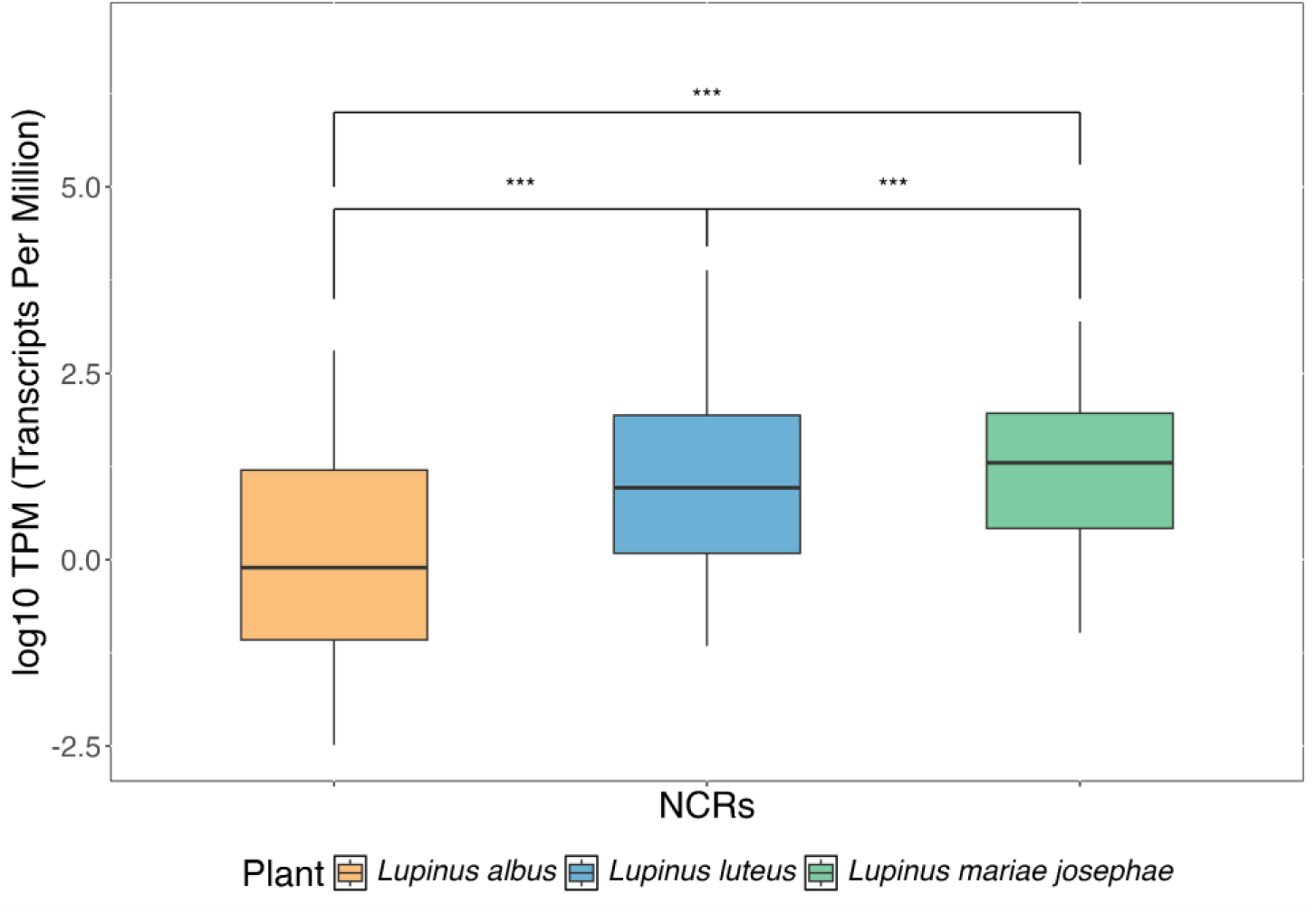
Expression of NCR genes across three *Lupinus* species. Boxplots show the distribution of log10-transformed TPM (Transcripts Per Million) values for nodule-specific NCR (Nodule Cysteine-Rich) genes in *Lupinus albus* (orange), *Lupinus luteus* (blue), and *Lupinus mariae-josephae* (green). Each box represents the interquartile range (IQR), with the horizontal line indicating the median. Whiskers extend to the furthest data points within 1.5×IQR from the lower and upper quartiles. Statistical significance was assessed using pairwise Wilcoxon rank-sum tests, and asterisks indicate highly significant differences (***: p < 0.001).

**Figure S6.**
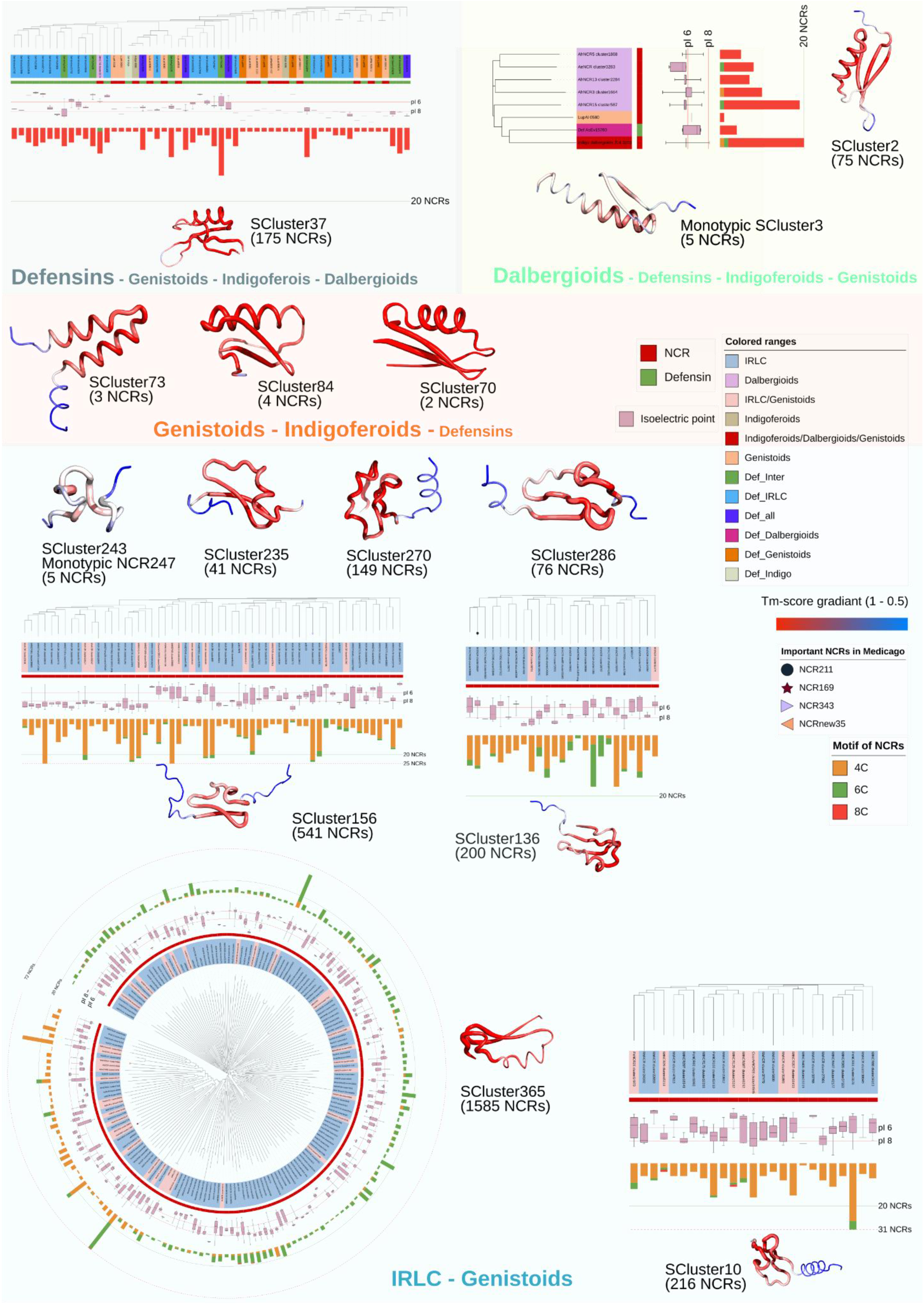
Structural phylogenetic analysis of each structural supercluster across legumes. Structure-based Foldtree phylogenetic trees of the most representative superclusters and structural alignments of these and other small superclusters. In each tree, the labels are colored according to the clade. The box plots represent the isoelectric point distributions. The multibar plots represent the size of clusters (number of NCR peptides), where the red represents the number of NCR peptides with an 8C motif, green with a 6C motif, and orange NCRs with a 4C motif. The structural alignments represented by a sausage representation are the alignments of all the structures of the supercluster with a color gradient from red (TM score 1) to blue (TM score 0.5). The size of the clades’ words inside the superclusters depends on the number of NCR peptides from this clade in the supercluster. The identified NCR peptides essential for an effective symbiosis in *M. truncatula* (if present in the presented superclusters) are annotated in the branches with different motifs.

**Figure S7.**
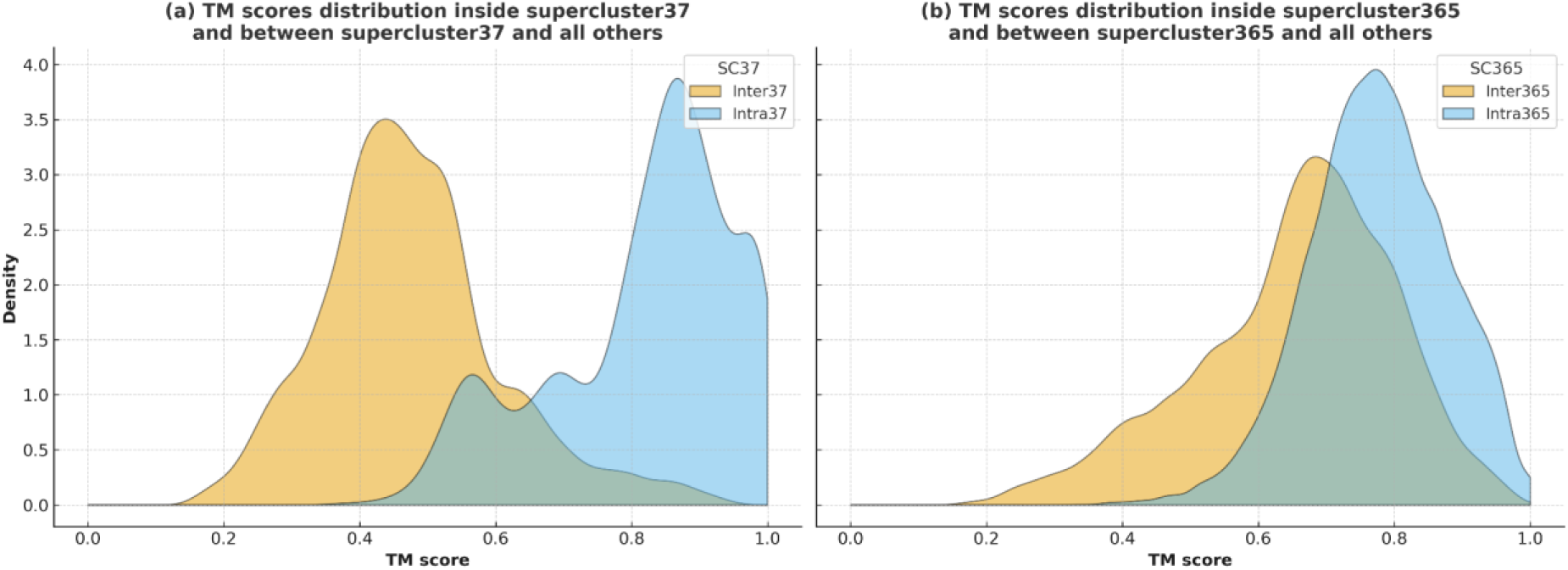
IRLC and Genistoids superclusters are conserved, while Dalbergioids superclusters are highly divergent. Comparison between TM scores inside one supercluster (blue) and between one supercluster and all the others (yellow), where two different profiles are observed. **(a)** The first is when we compare Dalbergioids (or defensins) with all other superclusters (IRLC-Genistoids), where the TM scores, in this case, are low for the inter-supercluster comparisons (density yellow up plots). **(b)** The second is when we compare IRLC-Genistoids superclusters with all others (other IRLC-Genisoids and only two superclusters of defensins-Dalbergioids), in which case the TM scores are high for the inter-supercluster comparisons (almost overlapping the TM scores inside the supercluster), indicating the conservation and homology of IRLC and Genistoids NCRs even across superclusters.

**Figure S8.**
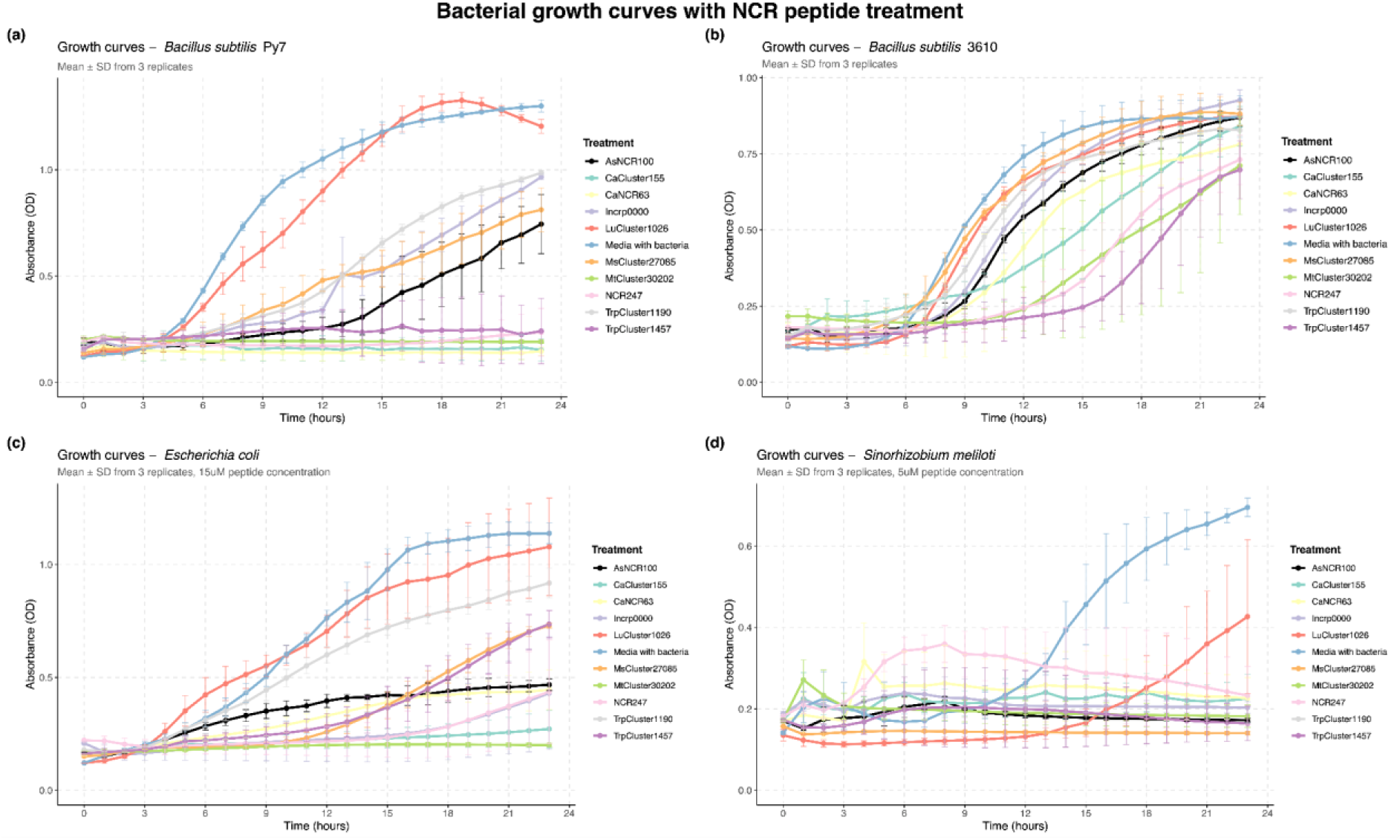
Growth curves of bacterial strains treated with NCR peptides and untreated control. **(a)** *Bacillus subtilis* strain Py7 growth curves over 24 hours following treatment with two µM NCR peptides. Bacterial growth was computed by optical density at 600 nm (OD₆₀₀). The untreated control is shown in light blue, with different NCR peptides, and the control peptide NCR247 is displayed in light pink. **(b)** *Bacillus subtilis* strain 3610 growth curves under identical treatment conditions as panel (a). **(c)** *Escherichia coli* growth curves following treatment with 15 µM NCR peptides and control bacteria. The higher peptide concentration accounted for the outer membrane permeability barrier characteristic of Gram-negative bacteria. **(d)** *Sinorhizobium meliloti* growth curves with five µM peptide treatments. *S. meliloti* is a nitrogen-fixing symbiotic bacterium that naturally encounters NCR peptides within legume root nodules. All data points represent mean ± standard deviation from three independent biological replicates. Error bars indicate standard deviation. Bacterial growth was computed hourly for 24 hours. Cultures were grown in LB medium (*B. subtilis* and *E. coli*) or TY medium (*S. meliloti*) with continuous shaking at 200 rpm. Peptides were added at time point 0 to exponentially growing cultures with an initial OD₆₀₀ of approximately 0.1. The different peptide concentrations (2 µM, five µM, and 15 µM) were selected based on preliminary dose-response experiments to achieve measurable effects within the physiological range for each bacterial species.

**Figure S9.**
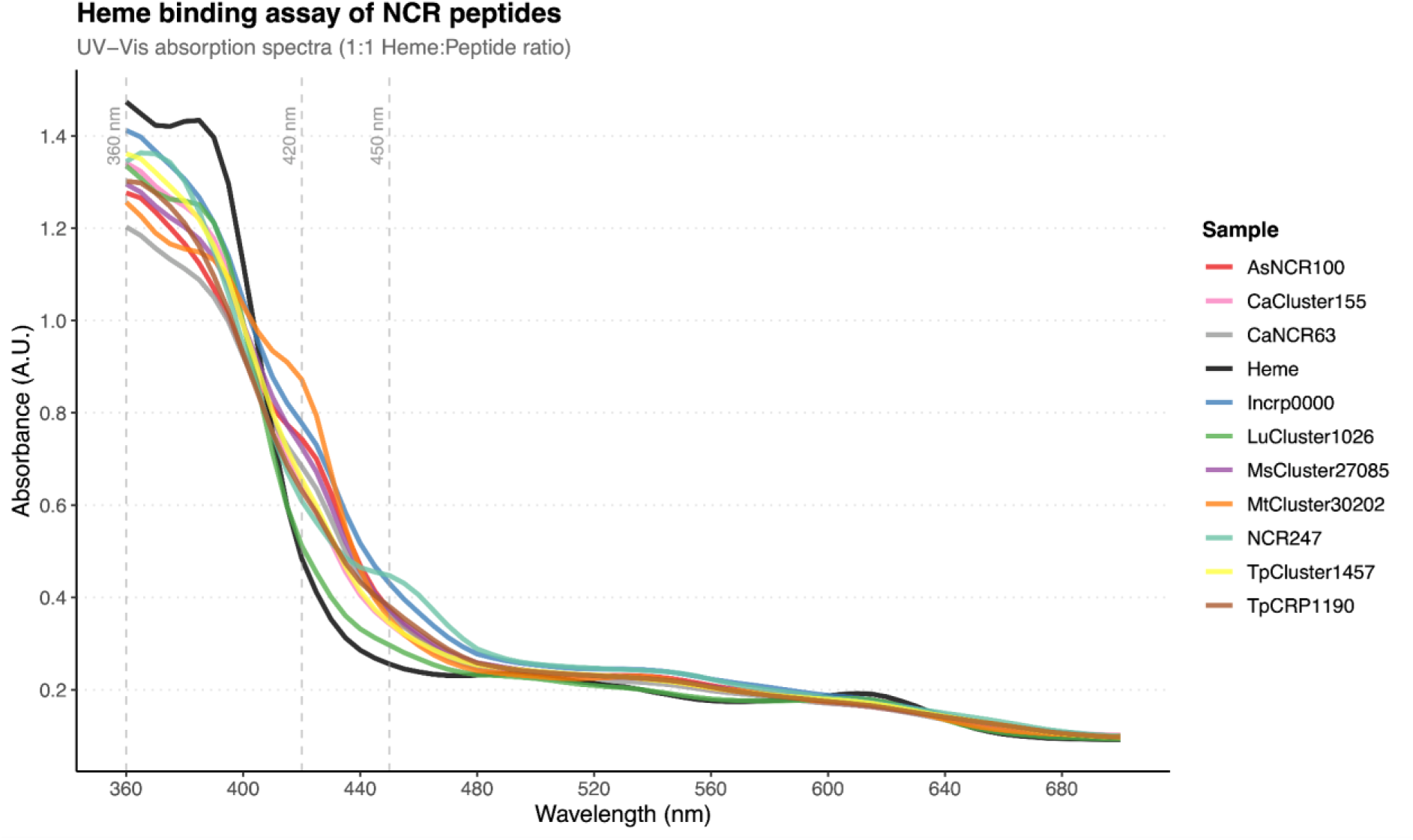
Heme binding assay of NCR peptides assessed by UV-Vis absorption spectroscopy. UV-Vis absorption spectra of heme alone (black line) and heme-NCR peptide complexes (colored lines) at a 1:1 molar ratio in phosphate buffer (pH 7.4) at room temperature. Measurements were performed from 360 to 700 nm with 5 nm intervals. Vertical dashed lines indicate key wavelengths for heme binding assessment: 360 nm and 450 nm (characteristic of NCR247-heme complex formation as previously reported), and 420 nm.

**Figure S10.**
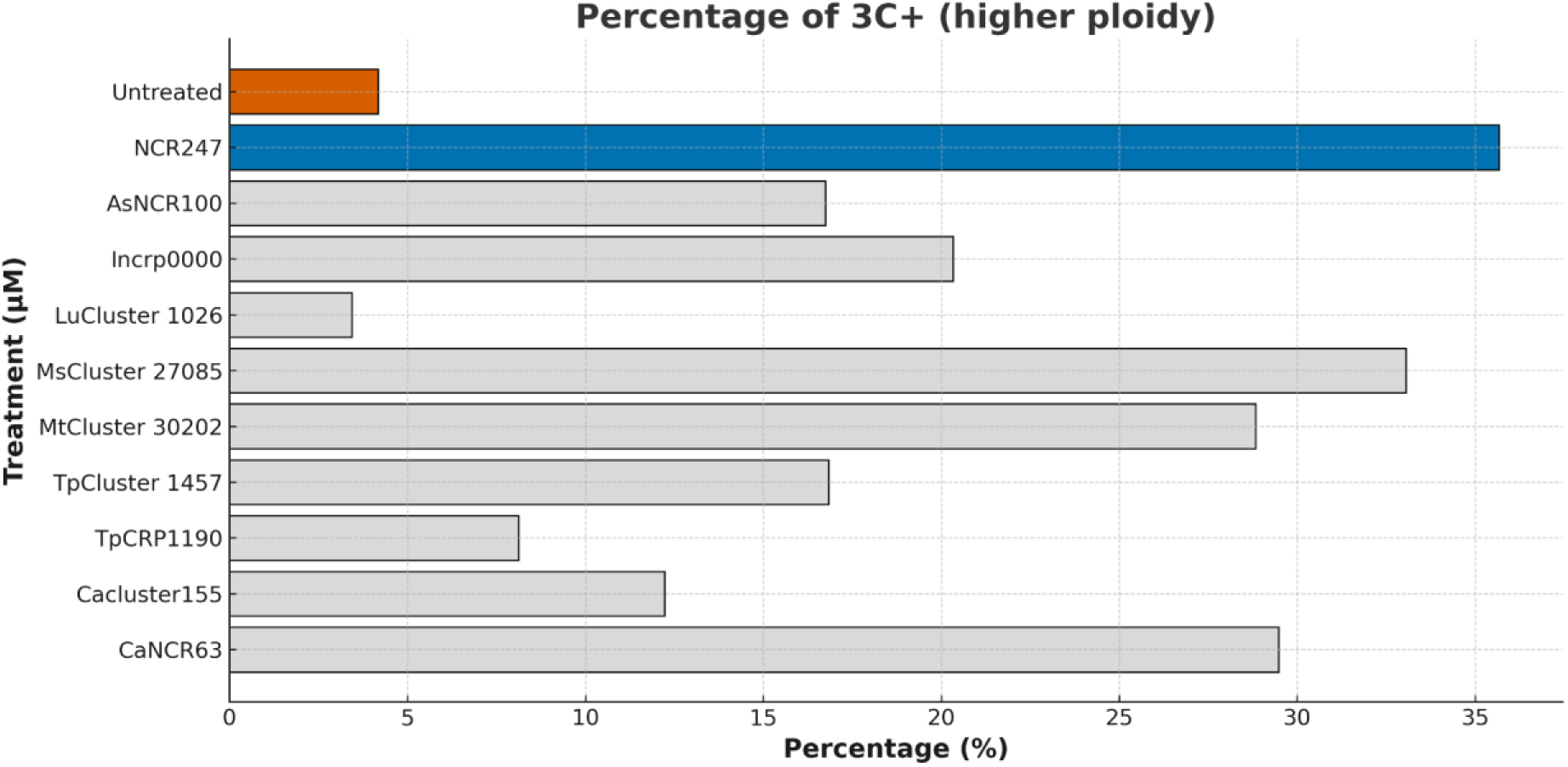
NCR peptides induce polyploidy (≥3C) in *S. meliloti* cells after 210 minutes of treatment with four μM. Bar plot showing the percentage of cells with ≥3C ploidy following treatment with various NCR peptides (4 µM) compared to untreated control. Each bar represents a different NCR peptide treatment.

## REFERENCES

Altschul, S. F., Gish, W., Miller, W., Myers, E. W., & Lipman, D. J. (1990). Basic local alignment search tool. Journal of Molecular Biology, 215(3), 403–410. 10.1016/S0022-2836(05)80360-2

Alunni, B., & Gourion, B. (2016). Terminal bacteroid differentiation in the legume−rhizobium symbiosis: Nodule-specific cysteine-rich peptides and beyond. New Phytologist, 211(2), 411–417. 10.1111/nph.14025

Alves-Carvalho, S., Aubert, G., Carrère, S., Cruaud, C., Brochot, A.-L., Jacquin, F., Klein, A., Martin, C., Boucherot, K., Kreplak, J., da Silva, C., Moreau, S., Gamas, P., Wincker, P., Gouzy, J., & Burstin, J. (2015). Full-length de novo assembly of RNA-seq data in pea (Pisum sativum L.) provides a gene expression atlas and gives insights into root nodulation in this species. The Plant Journal, 84(1), 1–19. 10.1111/tpj.12967

Andrews, S. (2010). FastQC: a quality control tool for high throughput sequence data. Babraham Bioinformatics, Babraham Institute, Cambridge, United Kingdom.

Audain, E., Ramos, Y., Hermjakob, H., Flower, D. R., & Perez-Riverol, Y. (2016). Accurate estimation of isoelectric point of protein and peptide based on amino acid sequences. Bioinformatics, 32(6), 821–827. 10.1093/bioinformatics/btv674

Barrière, Q., Guefrachi, I., Gully, D., Lamouche, F., Pierre, O., Fardoux, J., Chaintreuil, C., Alunni, B., Timchenko, T., Giraud, E., & Mergaert, P. (2017). Integrated roles of BclA and DD-carboxypeptidase 1 in Bradyrhizobium differentiation within NCR-producing and NCR-lacking root nodules. Scientific Reports, 7(1), Article 1. 10.1038/s41598-017-08830-0

Berendzen, J., Brown, A. V., Cameron, C. T., Campbell, J. D., Cleary, A. M., Dash, S., Hokin, S., Huang, W., Kalberer, S. R., Nelson, R. T., Redsun, S., Weeks, N. T., Wilkey, A., Farmer, A. D., & Cannon, S. B. (2021). The legume information system and associated online genomic resources. Legume Science, 3(3), e74. 10.1002/leg3.74

Camacho, C., Coulouris, G., Avagyan, V., Ma, N., Papadopoulos, J., Bealer, K., & Madden, T. L. (2009). BLAST+: Architecture and applications. BMC Bioinformatics, 10(1), 421. 10.1186/1471-2105-10-421

Chen, S. (2023). Ultrafast one-pass FASTQ data preprocessing, quality control, and deduplication using fastp. iMeta, 2(2), e107. 10.1002/imt2.107

Clevenger, J., Chu, Y., Scheffler, B., & Ozias-Akins, P. (2016). A Developmental Transcriptome Map for Allotetraploid Arachis hypogaea. Frontiers in Plant Science, 7. 10.3389/fpls.2016.01446

Collins, T. J. (2007). ImageJ for Microscopy. BioTechniques, 43(sup1), S25–S30. 10.2144/000112517

Czernic, P., Gully, D., Cartieaux, F., Moulin, L., Guefrachi, I., Patrel, D., Pierre, O., Fardoux, J., Chaintreuil, C., Nguyen, P., Gressent, F., Da Silva, C., Poulain, J., Wincker, P., Rofidal, V., Hem, S., Barrière, Q., Arrighi, J.-F., Mergaert, P., & Giraud, E. (2015). Convergent Evolution of Endosymbiont Differentiation in Dalbergioid and Inverted Repeat-Lacking Clade Legumes Mediated by Nodule-Specific Cysteine-Rich Peptides. Plant Physiology, 169(2), 1254–1265. 10.1104/pp.15.00584

Danecek, P., Bonfield, J. K., Liddle, J., Marshall, J., Ohan, V., Pollard, M. O., Whitwham, A., Keane, T., McCarthy, S. A., Davies, R. M., & Li, H. (2021). Twelve years of SAMtools and BCFtools. GigaScience, 10(2), giab008. 10.1093/gigascience/giab008

Davidson, N. M., Hawkins, A. D. K., & Oshlack, A. (2017). SuperTranscripts: A data driven reference for analysis and visualisation of transcriptomes. Genome Biology, 18(1), 148. 10.1186/s13059-017-1284-1

Davidson, N. M., & Oshlack, A. (2014). Corset: Enabling differential gene expression analysis for de novoassembled transcriptomes. Genome Biology, 15(7), 410. 10.1186/s13059-014-0410-6

Dobin, A., Davis, C. A., Schlesinger, F., Drenkow, J., Zaleski, C., Jha, S., Batut, P., Chaisson, M., & Gingeras, T. R. (2013). STAR: Ultrafast universal RNA-seq aligner. Bioinformatics, 29(1), 15–21. 10.1093/bioinformatics/bts635

Downie, J. A., & Kondorosi, E. (2021). Why Should Nodule Cysteine-Rich (NCR) Peptides Be Absent From Nodules of Some Groups of Legumes but Essential for Symbiotic N-Fixation in Others? Frontiers in Agronomy, 3. https://www.frontiersin.org/articles/10.3389/fagro.2021.654576

Eddy, S. R. (1998). Profile hidden Markov models. Bioinformatics, 14(9), 755–763. 10.1093/bioinformatics/14.9.755

Ekseth, O. K., Kuiper, M., & Mironov, V. (2014). orthAgogue: An agile tool for the rapid prediction of orthology relations. Bioinformatics, 30(5), 734–736. 10.1093/bioinformatics/btt582

Ellis, B., Haaland, P., Hahne, F., Le Meur, N., Gopalakrishnan, N., Spidlen, J., Jiang, M., & Finak, G. (2019). flowCore: Basic structures for flow cytometry data. R Package Version, 1(0).

Enright, A. J., Van Dongen, S., & Ouzounis, C. A. (2002). An efficient algorithm for large-scale detection of protein families. Nucleic Acids Research, 30(7), 1575–1584. 10.1093/nar/30.7.1575

Farkas, A., Maróti, G., Dürgő, H., Györgypál, Z., Lima, R. M., Medzihradszky, K. F., Kereszt, A., Mergaert, P., & Kondorosi, É. (2014). Medicago truncatula symbiotic peptide NCR247 contributes to bacteroid differentiation through multiple mechanisms. Proceedings of the National Academy of Sciences, 111(14), 5183–5188. 10.1073/pnas.1404169111

Frickey, T., & Lupas, A. (2004). CLANS: A Java application for visualizing protein families based on pairwise similarity. Bioinformatics, 20(18), 3702–3704. 10.1093/bioinformatics/bth444

Fu, L., Niu, B., Zhu, Z., Wu, S., & Li, W. (2012). CD-HIT: Accelerated for clustering the next-generation sequencing data. Bioinformatics, 28(23), 3150–3152. 10.1093/bioinformatics/bts565

Gage, D. J. (2004). Infection and Invasion of Roots by Symbiotic, Nitrogen-Fixing Rhizobia during Nodulation of Temperate Legumes. Microbiology and Molecular Biology Reviews, 68(2), 280–300. 10.1128/mmbr.68.2.280-300.2004

Gao, F., Yang, J., Zhai, N., Zhang, C., Ren, X., Zeng, Y., Chen, Y., Chen, R., & Pan, H. (2023). NCR343 is required to maintain the viability of differentiated bacteroids in nodule cells in Medicago truncatula. New Phytologist, 240(2), 815–829. 10.1111/nph.19180

Glazebrook, J., Ichige, A., & Walker, G. C. (1993). A Rhizobium meliloti homolog of the Escherichia coli peptide-antibiotic transport protein SbmA is essential for bacteroid development. Genes & Development, 7(8), 1485–1497. 10.1101/gad.7.8.1485

Grabherr, M. G., Haas, B. J., Yassour, M., Levin, J. Z., Thompson, D. A., Amit, I., Adiconis, X., Fan, L., Raychowdhury, R., Zeng, Q., Chen, Z., Mauceli, E., Hacohen, N., Gnirke, A., Rhind, N., di Palma, F., Birren, B. W., Nusbaum, C., Lindblad-Toh, K., … Regev, A. (2011). Full-length transcriptome assembly from RNA-Seq data without a reference genome. Nature Biotechnology, 29(7), Article 7. 10.1038/nbt.1883

Guefrachi, I., Nagymihaly, M., Pislariu, C. I., Van de Velde, W., Ratet, P., Mars, M., Udvardi, M. K., Kondorosi, E., Mergaert, P., & Alunni, B. (2014). Extreme specificity of NCR gene expression in Medicago truncatula. BMC Genomics, 15(1), 712. 10.1186/1471-2164-15-712

Guefrachi, I., Pierre, O., Timchenko, T., Alunni, B., Barrière, Q., Czernic, P., Villaécija-Aguilar, J.-A., Verly, C., Bourge, M., Fardoux, J., Mars, M., Kondorosi, E., Giraud, E., & Mergaert, P. (2015). Bradyrhizobium BclA Is a Peptide Transporter Required for Bacterial Differentiation in Symbiosis with Aeschynomene Legumes. Molecular Plant-Microbe Interactions®, 28(11), 1155–1166. 10.1094/MPMI-04-15-0094-R

Guerra-Garcia, F. J., & Sankari, S. (2025). NCR peptides in plant–bacterial symbiosis: Applications and importance. Trends in Microbiology, 33(2), 147–150. 10.1016/j.tim.2024.11.012

Haag, A. F., Baloban, M., Sani, M., Kerscher, B., Pierre, O., Farkas, A., Longhi, R., Boncompagni, E., Hérouart, D., Dall’Angelo, S., Kondorosi, E., Zanda, M., Mergaert, P., & Ferguson, G. P. (2011). Protection of Sinorhizobium against Host Cysteine-Rich Antimicrobial Peptides Is Critical for Symbiosis. PLOS Biology, 9(10), e1001169. 10.1371/journal.pbio.1001169

Haag, A. F., Kerscher, B., Dall’Angelo, S., Sani, M., Longhi, R., Baloban, M., Wilson, H. M., Mergaert, P., Zanda, M., & Ferguson, G. P. (2012). Role of Cysteine Residues and Disulfide Bonds in the Activity of a Legume Root Nodule-specific, Cysteine-rich Peptide*. Journal of Biological Chemistry, 287(14), 10791–10798. 10.1074/jbc.M111.311316

Haag, A. F., & Mergaert, P. (2020). Terminal bacteroid differentiation in the Medicago– Rhizobium interaction – a tug of war between plant and bacteria. In The Model Legume Medicago truncatula (pp. 600–616). John Wiley & Sons, Ltd. 10.1002/9781119409144.ch75

Horváth, B., Domonkos, Á., Kereszt, A., Szűcs, A., Ábrahám, E., Ayaydin, F., Bóka, K., Chen, Y., Chen, R., Murray, J. D., Udvardi, M. K., Kondorosi, É., & Kaló, P. (2015). Loss of the nodule-specific cysteine rich peptide, NCR169, abolishes symbiotic nitrogen fixation in the Medicago truncatula dnf7 mutant. Proceedings of the National Academy of Sciences, 112(49), 15232–15237. 10.1073/pnas.1500777112

Horváth, B., Güngör, B., Tóth, M., Domonkos, Á., Ayaydin, F., Saifi, F., Chen, Y., Biró, J. B., Bourge, M., Szabó, Z., Tóth, Z., Chen, R., & Kaló, P. (2023). The Medicago truncatula nodule-specific cysteine-rich peptides, NCR343 and NCR-new35 are required for the maintenance of rhizobia in nitrogen-fixing nodule*s* (p. 2023.01.23.523609). bioRxiv. 10.1101/2023.01.23.523609

Howe, K., Bateman, A., & Durbin, R. (2002). QuickTree: Building huge Neighbour-Joining trees of protein sequences. Bioinformatics, 18(11), 1546–1547. 10.1093/bioinformatics/18.11.1546

Huang, R., Snedden, W. A., & diCenzo, G. C. (2022). Reference nodule transcriptomes for Melilotus officinalis and Medicago sativa cv. Algonquin. Plant Direct, 6(6), e408. 10.1002/pld3.408

Illergård, K., Ardell, D. H., & Elofsson, A. (2009). Structure is three to ten times more conserved than sequence—A study of structural response in protein cores. Proteins, 77(3), 499–508. 10.1002/prot.22458

Jenei, S., Tiricz, H., Szolomájer, J., Tímár, E., Klement, É., Al Bouni, M. A., Lima, R. M., Kata, D., Harmati, M., Buzás, K., Földesi, I., Tóth, G. K., Endre, G., & Kondorosi, É. (2020). Potent Chimeric Antimicrobial Derivatives of the Medicago truncatula NCR247 Symbiotic Peptide. Frontiers in Microbiology, 11. 10.3389/fmicb.2020.00270

Johnson, L. S., Eddy, S. R., & Portugaly, E. (2010). Hidden Markov model speed heuristic and iterative HMM search procedure. BMC Bioinformatics, 11, 431. 10.1186/1471-2105-11-431

Jumper, J., Evans, R., Pritzel, A., Green, T., Figurnov, M., Ronneberger, O., Tunyasuvunakool, K., Bates, R., Žídek, A., Potapenko, A., Bridgland, A., Meyer, C., Kohl, S. A. A., Ballard, A. J., Cowie, A., Romera-Paredes, B., Nikolov, S., Jain, R., Adler, J., … Hassabis, D. (2021). Highly accurate protein structure prediction with AlphaFold. Nature, 596(7873), 583–589. 10.1038/s41586-021-03819-2

Kant, C., Pradhan, S., & Bhatia, S. (2016). Dissecting the Root Nodule Transcriptome of Chickpea (Cicer arietinum L.). PLOS ONE, 11(6), e0157908. 10.1371/journal.pone.0157908

Kazmierczak, T., Nagymihály, M., Lamouche, F., Barrière, Q., Guefrachi, I., Alunni, B., Ouadghiri, M., Ibijbijen, J., Kondorosi, É., Mergaert, P., & Gruber, V. (2017). Specific Host-Responsive Associations Between Medicago truncatula Accessions and Sinorhizobium Strains. Molecular Plant-Microbe Interactions®, 30(5), 399–409. 10.1094/MPMI-01-17-0009-R

Keller, J., Imperial, J., Ruiz-Argüeso, T., Privet, K., Lima, O., Michon-Coudouel, S., Biget, M., Salmon, A., Aïnouche, A., & Cabello-Hurtado, F. (2018). RNA sequencing and analysis of three Lupinus nodulomes provide new insights into specific host-symbiont relationships with compatible and incompatible Bradyrhizobium strains. Plant Science, 266, 102–116. 10.1016/j.plantsci.2017.10.015

Kim, M., Chen, Y., Xi, J., Waters, C., Chen, R., & Wang, D. (2015). An antimicrobial peptide essential for bacterial survival in the nitrogen-fixing symbiosis. Proceedings of the National Academy of Sciences, 112(49), 15238–15243. 10.1073/pnas.1500123112

Kolde, R. (2010). pheatmap: Pretty Heatmaps (p. 1.0.13) [Dataset]. 10.32614/CRAN.package.pheatmap

Konagurthu, A. S., Whisstock, J. C., Stuckey, P. J., & Lesk, A. M. (2006). MUSTANG: A multiple structural alignment algorithm. Proteins: Structure, Function, and Bioinformatics, 64(3), 559–574. 10.1002/prot.20921

Kumar, S., Stecher, G., Suleski, M., & Hedges, S. B. (2017). TimeTree: A Resource for Timelines, Timetrees, and Divergence Times. Molecular Biology and Evolution, 34(7), 1812–1819. 10.1093/molbev/msx116

Lambert, I., Paysant-Le Roux, C., Colella, S., & Martin-Magniette, M.-L. (2020). DiCoExpress: A tool to process multifactorial RNAseq experiments from quality controls to co-expression analysis through differential analysis based on contrasts inside GLM models. Plant Methods, 16(1), 68. 10.1186/s13007-020-00611-7

Lamouche, F., Bonadé-Bottino, N., Mergaert, P., & Alunni, B. (2019). Symbiotic Efficiency of Spherical and Elongated Bacteroids in the Aeschynomene-Bradyrhizobium Symbiosis. Frontiers in Plant Science, 10, 377. 10.3389/fpls.2019.00377

Langmead, B., & Salzberg, S. L. (2012). Fast gapped-read alignment with Bowtie 2. Nature Methods, 9(4), Article 4. 10.1038/nmeth.1923

Lavin, M., Herendeen, P. S., & Wojciechowski, M. F. (2005). Evolutionary Rates Analysis of Leguminosae Implicates a Rapid Diversification of Lineages during the Tertiary. Systematic Biology, 54(4), 575–594. 10.1080/10635150590947131

Lê, S., Josse, J., & Husson, F. (2008). FactoMineR: An R Package for Multivariate Analysis. Journal of Statistical Software, 25, 1–18. 10.18637/jss.v025.i01

Letunic, I., & Bork, P. (2021). Interactive Tree Of Life (iTOL) v5: An online tool for phylogenetic tree display and annotation. Nucleic Acids Research, 49(W1), W293– W296. 10.1093/nar/gkab301

Love, M. I., Huber, W., & Anders, S. (2014). Moderated estimation of fold change and dispersion for RNA-seq data with DESeq2. Genome Biology, 15(12), 550. 10.1186/s13059-014-0550-8

Maróti, G., Downie, J. A., & Kondorosi, É. (2015). Plant cysteine-rich peptides that inhibit pathogen growth and control rhizobial differentiation in legume nodules. Current Opinion in Plant Biology, 26, 57–63. 10.1016/j.pbi.2015.05.031

Maróti, G., Kereszt, A., Kondorosi, É., & Mergaert, P. (2011). Natural roles of antimicrobial peptides in microbes, plants and animals. Research in Microbiology, 162(4), 363–374. 10.1016/j.resmic.2011.02.005

Martin, M. (2011). Cutadapt removes adapter sequences from high-throughput sequencing reads. EMBnet.Journal, 17(1), Article 1. 10.14806/ej.17.1.200

Mergaert, P., Nikovics, K., Kelemen, Z., Maunoury, N., Vaubert, D., Kondorosi, A., & Kondorosi, E. (2003). A Novel Family in Medicago truncatula Consisting of More Than 300 Nodule-Specific Genes Coding for Small, Secreted Polypeptides with Conserved Cysteine Motifs, Plant Physiology, 132(1), 161–173. 10.1104/pp.102.018192

Mergaert, P., Uchiumi, T., Alunni, B., Evanno, G., Cheron, A., Catrice, O., Mausset, A.-E., Barloy-Hubler, F., Galibert, F., Kondorosi, A., & Kondorosi, E. (2006). Eukaryotic control on bacterial cell cycle and differentiation in the Rhizobium–legume symbiosis. Proceedings of the National Academy of Sciences, 103(13), 5230–5235. 10.1073/pnas.0600912103

Moi, D., Bernard, C., Steinegger, M., Nevers, Y., Langleib, M., & Dessimoz, C. (2023). Structural phylogenetics unravels the evolutionary diversification of communication systems in gram-positive bacteria and their viruses (p. 2023.09.19.558401). bioRxiv. 10.1101/2023.09.19.558401

Montiel, J., Downie, J. A., Farkas, A., Bihari, P., Herczeg, R., Bálint, B., Mergaert, P., Kereszt, A., & Kondorosi, É. (2017). Morphotype of bacteroids in different legumes correlates with the number and type of symbiotic NCR peptides. Proceedings of the National Academy of Sciences, 114(19), 5041–5046. 10.1073/pnas.1704217114

Nallu, S., Silverstein, K. A. T., Zhou, P., Young, N. D., & VandenBosch, K. A. (2014). Patterns of divergence of a large family of nodule cysteine-rich peptides in accessions of Medicago truncatula. The Plant Journal, 78(4), 697–705. 10.1111/tpj.12506

Oldroyd, G. E. D. (2013). Speak, friend, and enter: Signalling systems that promote beneficial symbiotic associations in plants. Nature Reviews Microbiology, 11(4), Article 4. 10.1038/nrmicro2990

Oono, R., & Denison, R. F. (2010). Comparing Symbiotic Efficiency between Swollen versus Nonswollen Rhizobial Bacteroids. Plant Physiology, 154(3), 1541–1548. 10.1104/pp.110.163436

Oono, R., Schmitt, I., Sprent, J. I., & Denison, R. F. (2010). Multiple evolutionary origins of legume traits leading to extreme rhizobial differentiation. New Phytologist, 187(2), 508–520. 10.1111/j.1469-8137.2010.03261.x

O’Rourke, J. A., Iniguez, L. P., Fu, F., Bucciarelli, B., Miller, S. S., Jackson, S. A., McClean, P. E., Li, J., Dai, X., Zhao, P. X., Hernandez, G., & Vance, C. P. (2014). An RNA-Seq based gene expression atlas of the common bean. BMC Genomics, 15(1), 866. 10.1186/1471-2164-15-866

Ott, T., van Dongen, J. T., Gunther, C., Krusell, L., Desbrosses, G., Vigeolas, H., Bock, V., Czechowski, T., Geigenberger, P., & Udvardi, M. K. (2005). Symbiotic Leghemoglobins Are Crucial for Nitrogen Fixation in Legume Root Nodules but Not for General Plant Growth and Development. Current Biology, 15(6), 531–535. 10.1016/j.cub.2005.01.042

Pan, H., & Wang, D. (2017). Nodule cysteine-rich peptides maintain a working balance during nitrogen-fixing symbiosis. Nature Plants, 3(5), Article 5. 10.1038/nplants.2017.48

Patro, R., Duggal, G., Love, M. I., Irizarry, R. A., & Kingsford, C. (2017). Salmon provides fast and bias-aware quantification of transcript expression. Nature Methods, 14(4), Article 4. 10.1038/nmeth.4197

Pazhamala, L. T., Purohit, S., Saxena, R. K., Garg, V., Krishnamurthy, L., Verdier, J., & Varshney, R. K. (2017). Gene expression atlas of pigeonpea and its application to gain insights into genes associated with pollen fertility implicated in seed formation. Journal of Experimental Botany, 68(8), 2037–2054. 10.1093/jxb/erx010

Pecrix, Y., Staton, S. E., Sallet, E., Lelandais-Brière, C., Moreau, S., Carrère, S., Blein, T., Jardinaud, M.-F., Latrasse, D., Zouine, M., Zahm, M., Kreplak, J., Mayjonade, B., Satgé, C., Perez, M., Cauet, S., Marande, W., Chantry-Darmon, C., Lopez-Roques, C., … Gamas, P. (2018). Whole-genome landscape of Medicago truncatula symbiotic genes. Nature Plants, 4(12), 1017–1025. 10.1038/s41477-018-0286-7

Penterman, J., Abo, R. P., De Nisco, N. J., Arnold, M. F. F., Longhi, R., Zanda, M., & Walker, G. C. (2014). Host plant peptides elicit a transcriptional response to control the Sinorhizobium meliloti cell cycle during symbiosis. Proceedings of the National Academy of Sciences, 111(9), 3561–3566. 10.1073/pnas.1400450111

Petersen, T. N., Brunak, S., von Heijne, G., & Nielsen, H. (2011). SignalP 4.0: Discriminating signal peptides from transmembrane regions. Nature Methods, 8(10), Article 10. 10.1038/nmeth.1701

Putri, G. H., Anders, S., Pyl, P. T., Pimanda, J. E., & Zanini, F. (2022). Analysing high-throughput sequencing data in Python with HTSeq 2.0. Bioinformatics, 38(10), 2943– 2945. 10.1093/bioinformatics/btac166

Quilbé, J., Lamy, L., Brottier, L., Leleux, P., Fardoux, J., Rivallan, R., Benichou, T., Guyonnet, R., Becana, M., Villar, I., Garsmeur, O., Hufnagel, B., Delteil, A., Gully, D., Chaintreuil, C., Pervent, M., Cartieaux, F., Bourge, M., Valentin, N., … Arrighi, J.-F. (2021). Genetics of nodulation in Aeschynomene evenia uncovers mechanisms of the rhizobium–legume symbiosis. Nature Communications, 12(1), 829. 10.1038/s41467-021-21094-7

Raul, B., Bhattacharjee, O., Ghosh, A., Upadhyay, P., Tembhare, K., Singh, A., Shaheen, T., Ghosh, A. K., Torres-Jerez, I., Krom, N., Clevenger, J., Udvardi, M., E. Scheffler, B., Ozias Akins, P., Dutta Sharma, R., Bandyopadhyay, K., Gaur, V., Kumar, S., & Sinharoy, S. (2021). Microscopic and transcriptomic analyses of Dalbergoid legume peanut reveal a divergent evolution leading to Nod Factor dependent epidermal crack-entry and terminal bacteroid differentiation. Molecular Plant-Microbe Interactions®. 10.1094/MPMI-05-21-0122-R

Ren, G. (2018). The evolution of determinate and indeterminate nodules within the Papilionoideae subfamily. 10.18174/429101

Ribeiro, C. W., Baldacci-Cresp, F., Pierre, O., Larousse, M., Benyamina, S., Lambert, A., Hopkins, J., Castella, C., Cazareth, J., Alloing, G., Boncompagni, E., Couturier, J., Mergaert, P., Gamas, P., Rouhier, N., Montrichard, F., & Frendo, P. (2017). Regulation of Differentiation of Nitrogen-Fixing Bacteria by Microsymbiont Targeting of Plant Thioredoxin s1. Current Biology, 27(2), 250–256. 10.1016/j.cub.2016.11.013

Robinson, M. D., McCarthy, D. J., & Smyth, G. K. (2010). edgeR: A Bioconductor package for differential expression analysis of digital gene expression data. Bioinformatics, 26(1), 139–140. 10.1093/bioinformatics/btp616

Roy, P., Achom, M., Wilkinson, H., Lagunas, B., & Gifford, M. L. (2020). Symbiotic Outcome Modified by the Diversification from 7 to over 700 Nodule-Specific Cysteine-Rich Peptides. Genes, 11(4), Article 4. 10.3390/genes11040348

Saifi, F., Biró, J. B., Horváth, B., Vizler, C., Laczi, K., Rákhely, G., Kovács, S., Kang, M., Li, D., Chen, Y., Chen, R., Domonkos, Á., & Kaló, P. (2024). Two members of a Nodule-specific Cysteine-Rich (NCR) peptide gene cluster are required for differentiation of rhizobia in Medicago truncatula nodules. The Plant Journal, 119(3), 1508–1525. 10.1111/tpj.16871

Sakai, H., Naito, K., Ogiso-Tanaka, E., Takahashi, Y., Iseki, K., Muto, C., Satou, K., Teruya, K., Shiroma, A., Shimoji, M., Hirano, T., Itoh, T., Kaga, A., & Tomooka, N. (2015). The power of single molecule real-time sequencing technology in the de novo assembly of a eukaryotic genome. Scientific Reports, 5(1), 16780. 10.1038/srep16780

Salgado, M. G., Demina, I. V., Maity, P. J., Nagchowdhury, A., Caputo, A., Krol, E., Loderer, C., Muth, G., Becker, A., & Pawlowski, K. (2022). Legume NCRs and nodule-specific defensins of actinorhizal plants—Do they share a common origin? PLOS ONE, 17(8), e0268683. 10.1371/journal.pone.0268683

Sankari, S., Babu, V. M. P., Bian, K., Alhhazmi, A., Andorfer, M. C., Avalos, D. M., Smith, T. A., Yoon, K., Drennan, C. L., Yaffe, M. B., Lourido, S., & Walker, G. C. (2022). A haem-sequestering plant peptide promotes iron uptake in symbiotic bacteria. Nature Microbiology, 7(9), Article 9. 10.1038/s41564-022-01192-y

Shabab, M., Arnold, M. F. F., Penterman, J., Wommack, A. J., Bocker, H. T., Price, P. A., Griffitts, J. S., Nolan, E. M., & Walker, G. C. (2016). Disulfide cross-linking influences symbiotic activities of nodule peptide NCR247. Proceedings of the National Academy of Sciences, 113(36), 10157–10162. 10.1073/pnas.1610724113

Simão, F. A., Waterhouse, R. M., Ioannidis, P., Kriventseva, E. V., & Zdobnov, E. M. (2015). BUSCO: Assessing genome assembly and annotation completeness with single-copy orthologs. Bioinformatics, 31(19), 3210–3212. 10.1093/bioinformatics/btv351

Song, L., & Florea, L. (2015). Rcorrector: Efficient and accurate error correction for Illumina RNA-seq reads. GigaScience, 4(1), s13742–015-0089-y. 10.1186/s13742-015-0089-y

The UniProt Consortium. (2019). UniProt: A worldwide hub of protein knowledge. Nucleic Acids Research, 47(D1), D506–D515. 10.1093/nar/gky1049

TransDecoder/TransDecoder: TransDecoder source. (n.d.). Retrieved April 18, 2023, from https://github.com/TransDecoder/TransDecoder

Trujillo, D. I., Silverstein, K. A. T., & Young, N. D. (2019). Nodule-specific PLAT domain proteins are expanded in the Medicago lineage and required for nodulation. New Phytologist, 222(3), 1538–1550. 10.1111/nph.15697

Van de Velde, W., Zehirov, G., Szatmari, A., Debreczeny, M., Ishihara, H., Kevei, Z., Farkas, A., Mikulass, K., Nagy, A., Tiricz, H., Satiat-Jeunemaître, B., Alunni, B., Bourge, M., Kucho, K., Abe, M., Kereszt, A., Maroti, G., Uchiumi, T., Kondorosi, E., & Mergaert, P. (2010). Plant Peptides Govern Terminal Differentiation of Bacteria in Symbiosis. Science, 327(5969), 1122–1126. 10.1126/science.1184057

Van Dongen, S. (2008). Graph Clustering Via a Discrete Uncoupling Process. SIAM Journal on Matrix Analysis and Applications, 30(1), 121–141. 10.1137/040608635

van Kempen, M., Kim, S. S., Tumescheit, C., Mirdita, M., Lee, J., Gilchrist, C. L. M., Söding, J., & Steinegger, M. (2024). Fast and accurate protein structure search with Foldseek. Nature Biotechnology, 42(2), 243–246. 10.1038/s41587-023-01773-0

Vincent, J. M. (1970). A manual for the practical study of the root-nodule bacteria. A Manual for the Practical Study of the Root-Nodule Bacteria. https://www.cabdirect.org/cabdirect/abstract/19710700726

Wang, Q., Yang, S., Liu, J., Terecskei, K., Ábrahám, E., Gombár, A., Domonkos, Á., Szűcs, A., Körmöczi, P., Wang, T., Fodor, L., Mao, L., Fei, Z., Kondorosi, É., Kaló, P., Kereszt, A., & Zhu, H. (2017). Host-secreted antimicrobial peptide enforces symbiotic selectivity in Medicago truncatula. Proceedings of the National Academy of Sciences, 114(26), 6854–6859. 10.1073/pnas.1700715114

Xu, D., Yang, Y., Gong, D., Chen, X., Jin, K., Jiang, H., Yu, W., Li, J., Zhang, J., & Pan, W. (2023). GFAP: Ultrafast and accurate gene functional annotation software for plants. Plant Physiology, 193(3), 1745–1748. 10.1093/plphys/kiad393

Yang, S., Wang, Q., Fedorova, E., Liu, J., Qin, Q., Zheng, Q., Price, P. A., Pan, H., Wang, D., Griffitts, J. S., Bisseling, T., & Zhu, H. (2017). Microsymbiont discrimination mediated by a host-secreted peptide in Medicago truncatula. Proceedings of the National Academy of Sciences, 114(26), 6848–6853. 10.1073/pnas.1700460114

Young, N. D., Debellé, F., Oldroyd, G. E. D., Geurts, R., Cannon, S. B., Udvardi, M. K., Benedito, V. A., Mayer, K. F. X., Gouzy, J., Schoof, H., Van de Peer, Y., Proost, S., Cook, D. R., Meyers, B. C., Spannagl, M., Cheung, F., De Mita, S., Krishnakumar, V., Gundlach, H., … Roe, B. A. (2011). The Medicago genome provides insight into the evolution of rhizobial symbioses. Nature, 480(7378), Article 7378. 10.1038/nature10625

Yuan, S., Zhou, S., Feng, Y., Zhang, C., Huang, Y., Shan, Z., Chen, S., Guo, W., Yang, H., Yang, Z., Qiu, D., Chen, H., & Zhou, X. (2021). Identification of the Important Genes of Bradyrhizobium diazoefficiens 113-2 Involved in Soybean Nodule Development and Senescence. Frontiers in Microbiology, 12. 10.3389/fmicb.2021.754837

Zhang, C., Shine, M., Pyle, A. M., & Zhang, Y. (2022). US-align: Universal structure alignments of proteins, nucleic acids, and macromolecular complexes. Nature Methods, 19(9), 1109–1115. 10.1038/s41592-022-01585-1

Zhang, S., Wang, T., Lima, R. M., Pettkó-Szandtner, A., Kereszt, A., Downie, J. A., & Kondorosi, E. (2023). Widely conserved AHL transcription factors are essential for NCR gene expression and nodule development in Medicago. Nature Plants, 9(2), Article 2. 10.1038/s41477-022-01326-4

Zhang, Y., & Skolnick, J. (2005). TM-align: A protein structure alignment algorithm based on the TM-score. Nucleic Acids Research, 33(7), 2302–2309. 10.1093/nar/gki524

Zhou, P., Silverstein, K. A., Gao, L., Walton, J. D., Nallu, S., Guhlin, J., & Young, N. D. (2013). Detecting small plant peptides using SPADA (Small Peptide Alignment Discovery Application). BMC Bioinformatics, 14, 335. 10.1186/1471-2105-14-335

Zorin, E. A., Kliukova, M. S., Afonin, A. M., Gribchenko, E. S., Gordon, M. L., Sulima, A. S., Zhernakov, A. I., Kulaeva, O. A., Romanyuk, D. A., Kusakin, P. G., Tsyganova, A. V., Tsyganov, V. E., Tikhonovich, I. A., & Zhukov, V. A. (2022). A variable gene family encoding nodule-specific cysteine-rich peptides in pea (Pisum sativum L.). Frontiers in Plant Science, 13. https://www.frontiersin.org/articles/10.3389/fpls.2022.884726

